# Harnessing Synaptic Vesicle Release and Recycling Mechanism for Molecule Delivery to Neurons

**DOI:** 10.1101/2024.09.11.612569

**Authors:** Karen Kar Lye Yee, Junichi Kumamoto, Daijiro Inomata, Naoki Suzuki, Ryuhei Harada, Norihiro Yumoto

## Abstract

Neurodegenerative clinical trials often fail due to insufficient drug doses in reaching targeted cells and the unintended delivery to non-targeted cells. This study demonstrates an alternative neuron-selective drug delivery system, which utilizes the synaptic vesicle release and recycling mechanism (SVRM) by antibody shuttles targeting synaptic vesicle transmembrane proteins for molecule delivery. Using Synaptotagmin-2 (SYT2), we exemplify that intravenously administered anti-SYT2 antibodies localize to neuromuscular junctions, undergo uptake, and retrograde transport to *ChAT-*positive motor neurons (MNs) in the spinal cord and brainstem. The delivery of anti-microtubule agent and *Malat1* gapmer antisense oligonucleotide to MNs with anti-SYT2 antibodies induces axon degeneration and reduction of Malat1 RNA expression, respectively. This approach circumvents the blood-spinal cord barrier, enabling selective delivery of therapeutic molecules to neurons while minimizing effects in non-targeted cells. Thus harnessing SVRM presents a promising strategy for enhancing drug concentrations in neurons and improving treatment efficacy for neurodegenerative diseases.

## Introduction

Developing effective therapies for neurodegenerative diseases remains one of the most challenging and urgent objectives in biomedical research. Key obstacles in this pursuit include the multifactorial pathophysiology nature of these disorders, and the unique barriers at the central nervous system (CNS); the blood-brain barrier (BBB) and the blood-spinal cord barrier (BSCB).^1, 2^ These barriers are composed of specialized endothelial cells, astrocytes, pericytes and a basement membrane, which form a highly selective interface between the blood and the brain or spinal cord tissue, respectively.^3, 4^ While essential for maintaining CNS homeostasis, these barriers present a formidable hurdle in drug development. Estimates suggest that over 95% of small molecule drugs and nearly all large molecule therapeutics fail to cross the BBB and BSCB effectively.^5, 6^

Hence, overcoming these barriers are crucial for advancing new therapeutics for neurodegenerative diseases. Various mechanisms utilizing receptor-mediated transcytosis, adeno-associated virus and cell penetrating peptides have been explored to circumvent the BBB and BSCB for therapeutic molecule delivery to cerebrospinal tissues.^7–10^ Among these, receptor-mediated transcytosis with engineered antibodies targeting the transferrin receptor has shown some promise enabling therapeutic molecules transcytosis across the BBB.^11–13^ However, it lacks cell specificity within CNS tissues and a mechanism to transverse neuronal cell membrane post-BBB entry.

The therapeutic efficacy of neurological treatments hinges not only on successful drug delivery to CNS but also on precise drug targeting to specific cell types within the CNS. The absence of cell specific targeting may result in suboptimal drug concentrations in targeted cells or off-target effects on non-targeted cells.^14^ Drug efficacy can also be constrained by insufficient or absence of efficient intracellular delivery systems to deliver drugs into specific cellular compartments of the targeted cell.^15^ This is particularly relevant for neurodegenerative conditions such as Amyotrophic Lateral Sclerosis, Alzheimer’s disease, and Parkinson’s disease, where specific neuronal populations are affected. In order to address these challenges, we focused on harnessing the synaptic vesicle release and recycling mechanism (SVRM) and utilizing an engineered antibody shuttle against the synaptic vesicle (SV) transmembrane proteins of neurons for therapeutic molecule delivery. This approach provides a means to transport molecules into cells without inducing non-physiological activities, such as receptor-mediated endocytosis observed in typical antibody drug conjugate in oncology, and it ensures neuron selectivity post-BBB or -BSCB transversal since SVRM is a unique feature for neurons.^16^

In this study, synaptotagmin-2 (SYT2) was selected to illustrate monoclonal antibodies raised against luminal domains of SV transmembrane proteins are capable of targeted molecule delivery to neurons through SVRM. Our immunohistological analyses demonstrate that the intravenously administered monoclonal SYT2 antibody shuttle (mAb-SYT2) was selectively and efficiently taken up by motor neurons (MNs) at the neuromuscular junctions (NMJs) and retrogradely transported to the soma in the spinal cord and brainstem. Furthermore, the *in vitro* and *in vivo* data show that *MALAT1* gapmer antisense oligonucleotides (ASO) conjugated with mAb-SYT2 reduces *MALAT1* RNA expression in targeted cells. This indicates that payloads delivered by mAb-SYT2 were successfully released into MNs cytoplasm.

Here we propose that antibodies shuttle targeting luminal domain of SV transmembrane proteins can utilize SVRM as an alternative route to deliver therapeutic molecules to intended neuronal populations.

## Results

### LRRTM2-Coated Microbeads Induces Synaptic Vesicle Release and Recycling Mechanism in Human iPSC-Derived MNs

The conventional method of delivering therapeutic molecules to neurons is through intrathecal injection which causes adverse effects in patients.^17, 18^ Here we propose a different route to deliver molecules by utilizing SVRM occurring at the presynapses. In order to validate this concept, as well as to acquire high-affinity antibodies that can target the luminal domains of SV transmembrane proteins and be efficiently internalized; an *in vitro* induced-presynapse model was developed. The *in vitro* induced-presynapse model uses microbeads coated with Leucine-rich repeat transmembrane protein 2 (LRRTM2), a purified postsynaptic membrane protein, to initiate presynapse differentiation in human induced pluripotent stem cell (iPSC)-derived MNs. Previous studies have shown that overexpression of synaptogenic transmembrane proteins in fibroblasts and coating microbeads with fabricated clusters of synaptogenic extracellular domains (ECDs) proteins can induce pre- and post-synaptic differentiation in cultured neurons.^19^ LRRTM2 is a well characterized postsynaptic protein known to induce presynaptic differentiation in hippocampal and cortical neurons in mice.^20–24^

Our findings demonstrated that microbeads coated with LRRTM2 ECDs fused to human immunoglobulin G (IgG) Fc domain induced human iPSC-derived MNs to form synapsin-1 aggregates at microbeads contact sites. Whereas control human IgG-Fc protein coated microbeads had no synapsin-1 aggregation effect at microbeads (Figure 1A-C, Figure S1A). The differentiated presynapse had indications of increased acetylcholine secretion detected in the culture medium (data not shown). These data indicate that LRRTM2 induces presynaptic differentiation in human MNs.

**Figure 1.**
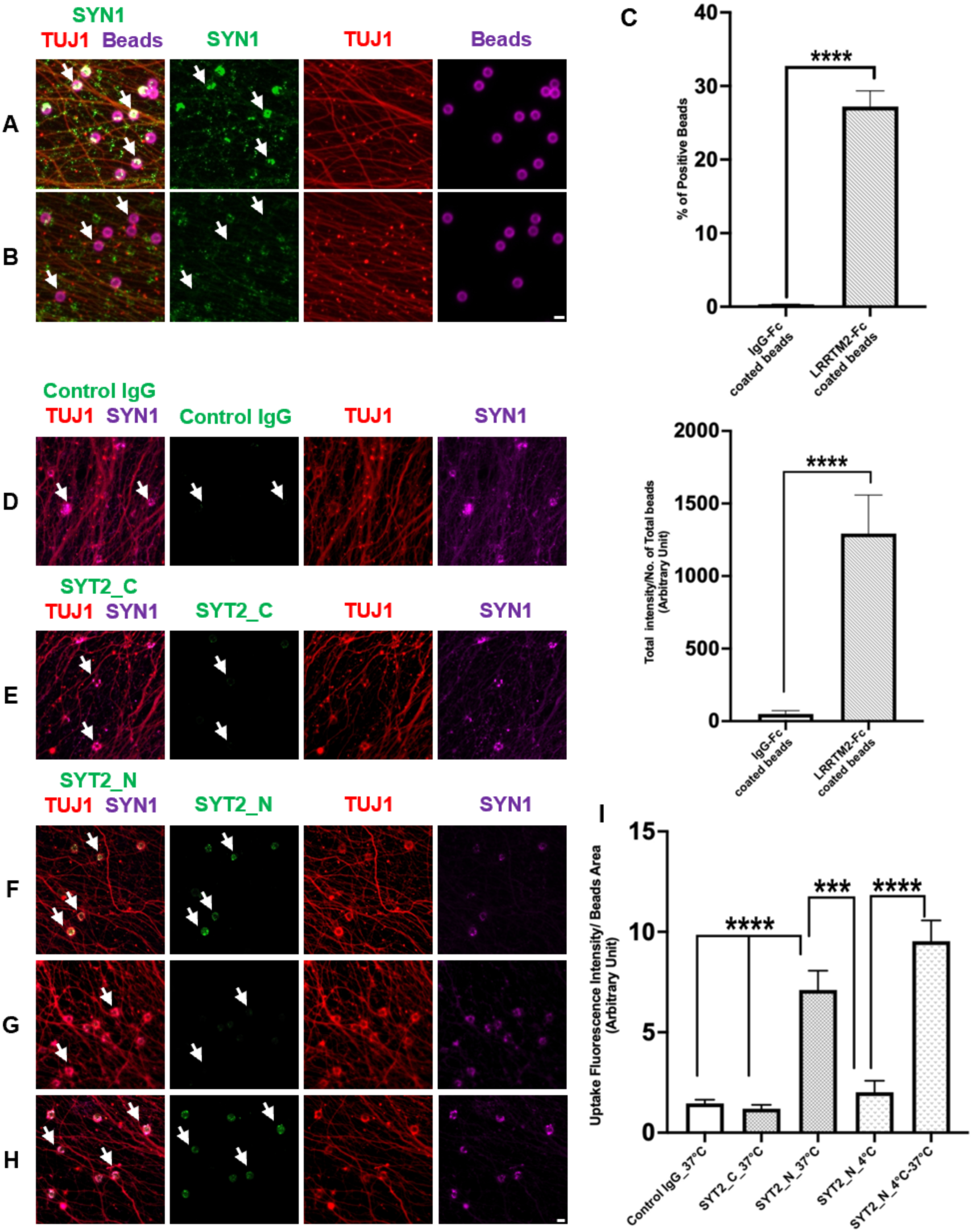
Induction of SVRM in human iPSC-derived MN (A) The addition of LRRTM2-coated microbeads induces presynapse differentiation (yellow, white arrows) with expression of SYN1 presynapse marker (green, white arrows), TUJ1 axon marker (red) and presence of LRRTM2-coated microbeads (magenta). (B) Minimal presynapse differentiation with control IgG-coated microbeads. (C) Quantification of presynapsed formed (top) and density of presynapse per bead (bottom), n=5. (F) Molecular shuttling through SVRM requires antibody specificity to the luminal domain of SV transmembrane protein. Luminal domain targeting antibody, SYT2 N-terminal, localizes to presynapse on LRRTM2-coated microbeads (yellow, white arrows). (D-E) Absence of presynapse localization with C-terminal antibodies targeting the cytoplasmic domain of SYT2 and control IgG (arrows). (G) Exo-endocytosis required for antibody shuttling into SVRM. At 4°C, there is a decrease in presynapse exocytosis activity with no localization of SYT2 N-terminal antibodies to the presynapse (arrows). (H) Recovery of exocytosis when reverted to 37°C. Localization of SYT2 N-terminal antibodies to the presynapse (green, white arrows) and merged images (yellow, white arrows). (I) Graphical representation of antibodies uptake at presynapse (D-H), n=4. Scale bars 10µm. All data expressed as mean±s.e.m; \*\*\*\**p*<0.0001, \*\*\**p*<0.001.

Furthermore, our immunostaining experiments on human muscle specimens revealed an accumulation of LRRTM2 proteins at NMJs, suggesting a functional role for LRRTM2 in formation and/or maintenance of human NMJs (Figure S1B).

Antibody shuttles using SVRM requires high selectivity and affinity for the luminal domain of SV transmembrane proteins, which are transiently exposed during presynapse SV exocytosis, are criteria for neuronal shuttle suitability. In order to identify that the luminal domain exposure is crucial for antibody binding followed by internalization into neurons, we used the *in vitro* induced-presynapse model and commercially available anti-SYT2 N-terminal (luminal domain) rabbit polyclonal antibody for antibody uptake study. Neuronal stimulation with 4-aminopyridine (4-AP) at 37°C showed SYT2 antibody internalization (Figure 1F). On the other hand, control normal rabbit IgG and anti-SYT2 C-terminal (cytoplasmic domain) rabbit polyclonal antibodies were not internalized when stimulated with 4-AP at 37°C. (Figure 1D-E). To demonstrate that the antibody uptake depends on SVRM and is not dependent on the presence of 4-AP, we investigated the temperature dependency required for antibody internalization. At 4°C, the reduction of exo-endocytotic activities inhibits SYT2 N-terminal antibody uptake (Figure 1G). When the iPSC-derived MNs were reverted to 37°C, anti-SYT2 N-terminal antibody localization to the SV was again observed (Figure 1H-I). This suggests the SVRM antibody shuttle ability to deliver therapeutic molecules to neurons, which is dependent on the neuronal exo-endocytosis process and a specificity for SV luminal domain, is feasible.

### Generation and Characteristics of Monoclonal Antibodies Targeting Luminal Domains of SV Transmembrane Protein

After conceptualizing SVRM as a potential route for antibody shuttle uptake, monoclonal antibodies that had high specificity for luminal domain of SV transmembrane protein were required. Several transmembrane proteins localized in the SV are involved in neurotransmission. Through published literature and public domain databases, 16 such potential candidates were identified. Of which, we selected SYT2 to exemplify our concept further. SYT2 being a single-pass membrane protein allows the ease of monoclonal antibody generation.^25^ Furthermore, its main physiological function in calcium sensing and SV recruitment mechanism to the active zone for neurotransmitter release is located at the cytoplasmic C-terminal region consisting of 336 amino acid residues. We postulate that generating antibodies against the luminal domain, full 62 and partial 25 amino acid residues, will have the minimum physiological hindrance. In addition, SYT2 accumulates at all NMJs whereas SYT1 with the closest homology to SYT2 among 17 synaptotagmin family members, accumulates at less than 50% of NMJs in mice and rats.^26, 27^ Thus, SYT2 monoclonal antibodies are described here while the rest of the generated antibodies against other SV proteins antigens are not shown (Figure S1C).

Based on FACS and ELISA screening, primary and secondary candidates were identified from 37 scFv clones against human SYT2 full N-terminal domain (h2FL,1-62 a.a.) and 93 clones against the partial N-terminal domain (h2PL,1-25 a.a.) were allocated into seven clusters each. These scFvs are reactive to both human SYT2 and mouse Syt2, but not to human SYT1 (data not shown). For final selection, we examined the binding and internalizing capabilities of these scFv clusters with the *in vitro* induced-presynapse model. Two h2FL and four h2PL were identified to efficiently internalized into presynaptic specializations induced by synaptogenic microbeads on human iPS-derived MNs in the presence of 4-AP (Figure S1D-E). The characteristics of these six monoclonal SYT2 scFvs are summarized in table S1. Selected anti-SYT2 scFvs were converted to chimeric full-length IgGs with their Fc regions are of human Fc regions (mAb-SYT2). Hereafter, all data shown in this report were collected by using chimeric mAb-SYT2 clone against full luminal domain (FL) and partial luminal domain (PL), unless otherwise described.

Dissociation constant (K_D_) of these six mAb-SYT2 was conducted. Since there was no commercially available monoclonal antibody which is reactive to human SYT2, we used K_D_ of FL01 converted to chimeric full-length IgG to benchmarked the six antibodies K_D_. Our data shows FL01 falls within the median cut off range in the candidate selection uptake study (Figure S1D). A ratio higher than FL01 indicates lesser SYT2 antigen affinity and a lower ratio indicates greater SYT2 antigen affinity (Table S2). PL13 showed a median ratio value of 1.7 and was selected for subsequent studies.

### *In vivo* Uptake and Distribution of SYT2 Antibody Shuttle

Neuronal uptake of PL13 upon intravenous (IV) injection at various time points in mice was investigated by immunostaining with fluorophore-conjugated anti-human IgG Fc domain antibodies. The PL13 signals were localized at NMJs, which were co-visualized with synapsin-1 antibody and α-bungarotoxin, from 12h-240h post-IV injection in gastrocnemius (GAS) and tibialis anterior (TA) muscle tissue (Figure 2A-B). Random localization of control IgG antibodies at NMJs was an undetected amount at all time points. The 240h data is shown in figure 2A-B, right. In GAS and TA, PL13 localization was observed at 12h post-IV injection. GAS maintained comparable intensity across all time points, suggesting continuous uptake, while TA uptake peaked at 72h post-IV injection (Figure 2A-B).

**Figure 2.**
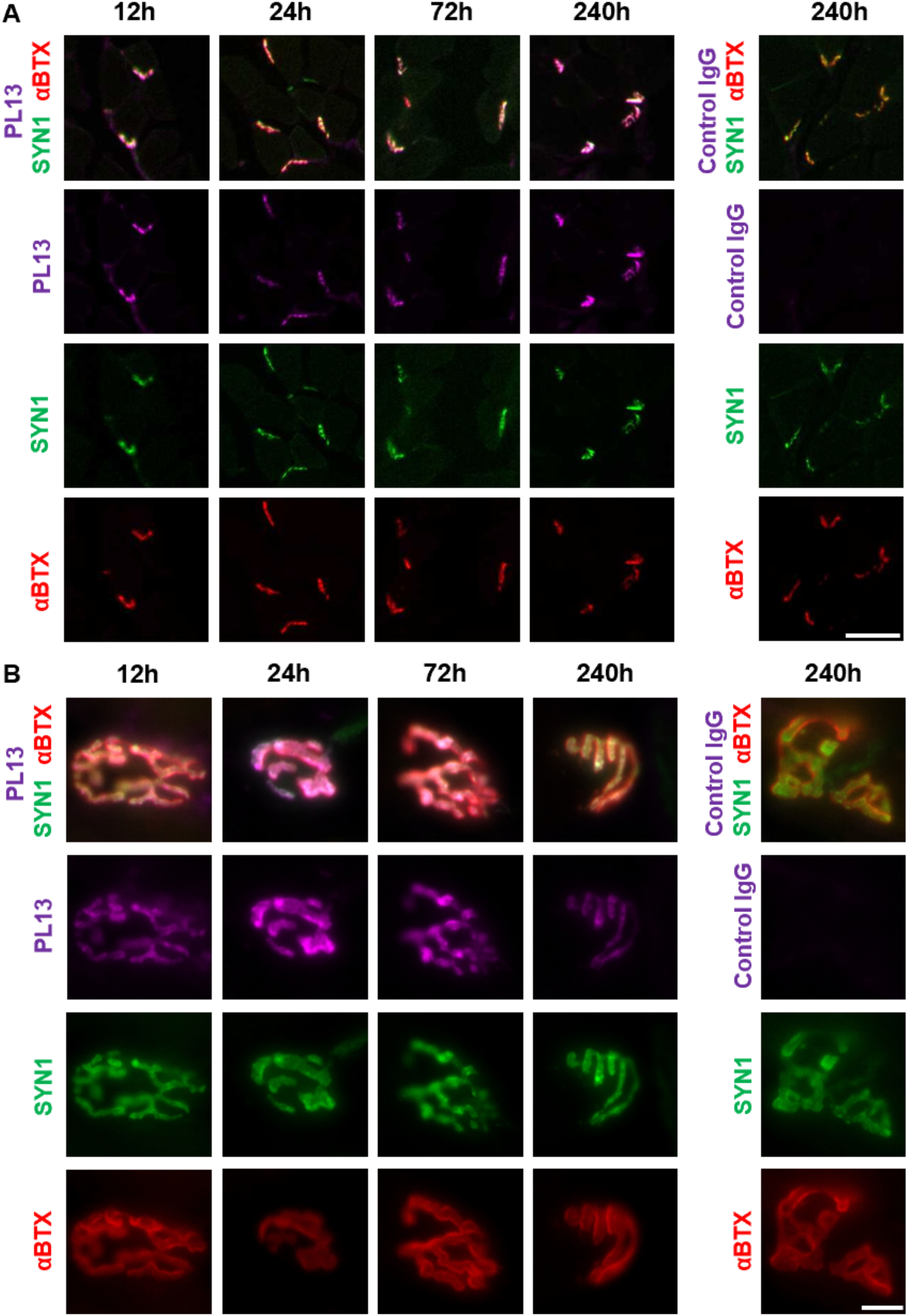
NMJ uptake of SVRM molecular shuttle post-intravenous administration (A) PL13 uptake at NMJ of GAS tissues were observed at 12h-240h (magenta) but not with control IgG up to 240h (right). Scale bar 50µm. (B) PL13 uptake at NMJ of TA tissues were also observed (magenta) from 12-240h but not with control IgG antibodies (right). Scale bar 10µm.

To maintain neuronal homeostasis, it is essential for damaged, aging proteins and organelles, including SVs, to be removed from the axon terminals and distal axon by retrograde transport to the soma for degradation and recycling of components. Hence, we investigated the distribution of PL13 antibodies to the soma. Frozen sections of the cervical spinal cords from the same animals used in NMJ localization experiments were co-immunostained with ChAT antibody and anti-human IgG Fc antibody in the soma. Weak signals of PL13 were observed at 12h-24h and accumulated intensely at 72h-240h in ChAT-positive MNs (Figure 3A, S2A). Control IgG antibodies in ChAT-positive MNs were undetected at all time points (Figure 3A, right). Interestingly, the subcellular localization of PL13 signals in the soma were co-stained in lysosome with LAMP1. This suggests that endosomes and/or autophagosomes generated from post-SVRM uptake of PL13 underwent cargo sorting and were transported to the soma for protein degradation (Figure 3B).

**Figure 3.**
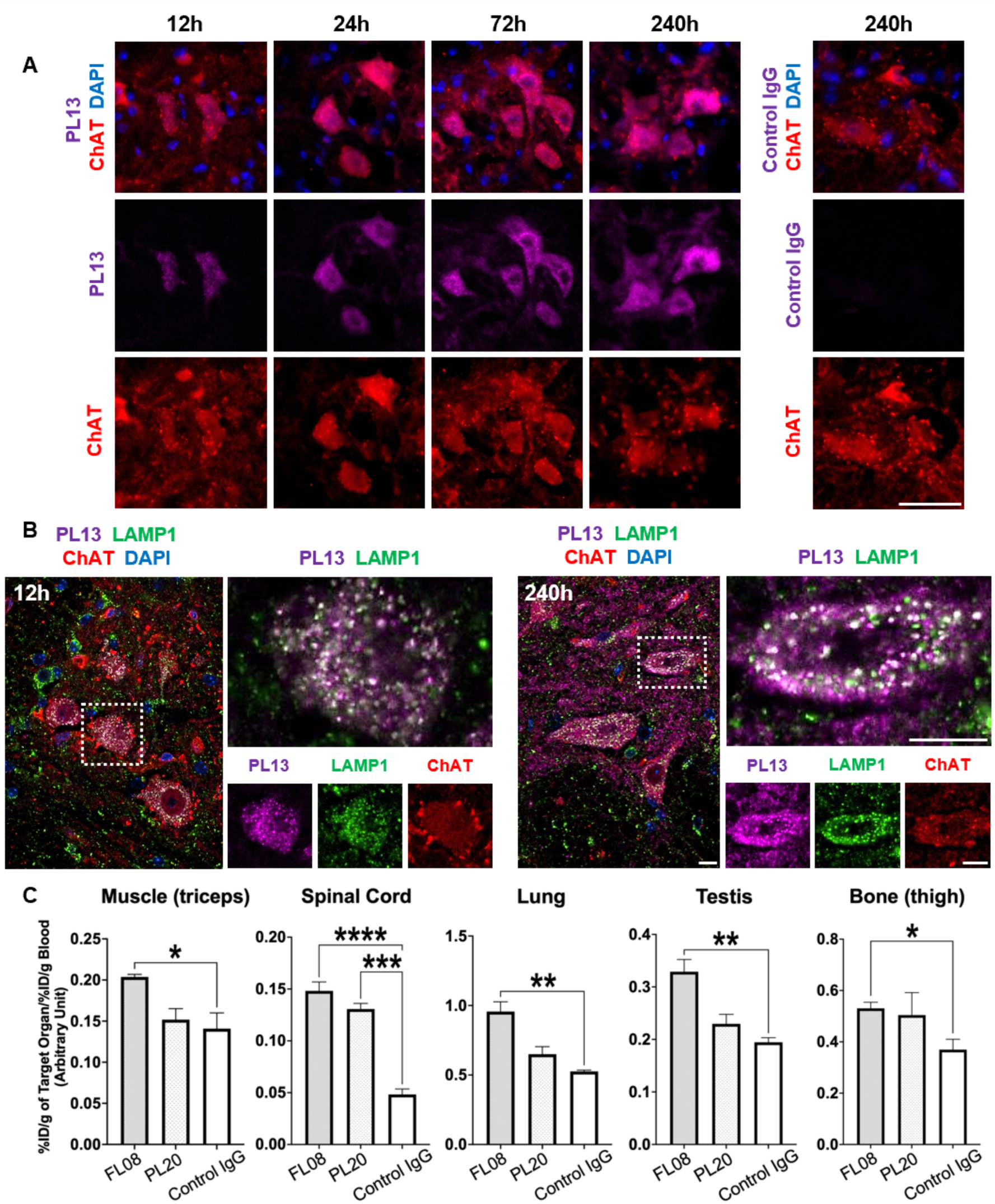
*in vivo* distribution of SVRM molecular shuttle post-intravenous administration. (A) 12h post-NMJ uptake of PL13 (magenta), antibodies retrograde to MN soma located in the ventral horn of the spinal cord. From 72h, PL13 was intensely localized to ChAT (red) stained MN. Scale bar 50µm. See also Figure S2a. (B) Detection of PL13 (magenta) in lysosomes (Lamp1, green) of MN soma from 12h and 240h. Indicates the shuttle is retrograded to the spinal cord through the endo-lysosome system after SVRM uptake. Scale bar 10µm. (C) FL08 and PL20 were conjugated with radioisotope Zr^89^. FL08-Zr^89^ significantly localizes to muscle, lung, bone and testis compared to control IgG-Zr^89^. Both FL08 and PL20 substantially retrogrades more Zr^89^ to the spinal cord compared to control IgG-Zr^89^. Data expressed as mean±s.e.m., n=3 and T-test analysis; \*\*\*\**p*<0.0001, \*\*\**p*<0.001, \*\**p*<0.01, \**p*<0.05. See also Figure S2B and Table S3.

In liver, PL13 and control IgG were detected at 12h-24h and by 72h both antibodies were not detectable in the liver (Figure S3A-B), This indicates that there is no difference in antibody clearance rate between mAb-SYT2 and control IgG in the liver. Most cranial MNs originate from the brainstem and innervate peripheral muscles in the head and neck region, controlling functions such as eye movement, facial expression, mastication, swallowing, and speech. Subsequently, PL13 uptake through SVRM and retrograde transport to the soma in the cranial MNs was examined. Day 10 post-IV, brainstem frozen sections were analyzed with immunostaining.

Visualization of PL13 revealed its localization in ChAT-positive somas of the oculomotor nucleus (3N), abducens nucleus (6N) and facial nucleus (7N) (Figure S3C). Intriguingly, the staining of intrinsically expressing SYT2 with rabbit polyclonal anti-SYT2 in the brainstem did not overlap with the PL13 signals. This indicates that PL13 after uptake localizes to regions different from intrinsic expressing SYT2 regions or there could be SYT2 competitive binding. Control IgG antibodies were not detected in the brainstem at all time points (data not shown). mAb-SYT2 shuttle distribution and uptake were further investigated with FL08 and PL20 conjugated with Zirconium radioisotope. At 72h, the uptake of mAb-SYT2, was quantified with a localization ratio which factors in blood clearance of the antibodies by determining the ratio of percentage of injected dose per gram (%ID/g) in organ and blood. The localization ratio showed FL08 was significantly detected in the lung, testis, muscle (triceps) and bone (thigh) compared with control IgG (Figure 3C).^28^

Both FL08 and PL20 exhibited significantly higher spinal cord localization ratios. In order to grasp the selectivity and efficacy of mAb-SYT2 tendency to retrograde to the spinal cord, we evaluated the immunospecificity index of a specific organ by dividing localization ratio of mAb-SYT2 against the control. Both FL08 and PL20 had a 3-2.5 fold immunospecificity index in the spinal cord post-IV injection (Table S3). This is noteworthy given that whole IgG antibodies have difficulty crossing the BSCB. The observed spinal cord penetration suggests antibodies shuttle utilizing SVRM properties at spinal motor neurons were able to achieve central nervous system entry.

### *In vitro* and *in vivo* Delivery of an Active Payload by mAb-SYT2

We next investigated mAb-SYT2 capability to deliver functional payloads into MNs in the *in vitro* induced-presynapse model. PL13 was conjugated with an anti-mitotic agent, monomethyl auristatin E (MMAE) and added to cultured MNs. Specific targeting of PL13 and 4-AP stimulation, functional MMAE was transported into MNs resulting in axon degeneration (Figure 4A, left). A decrease of 26.6% in axon area (11.67±1.096 s.e.m.) was observed compared with random uptake of control chimera IgG-MMAE (15.9±1.332 s.e.m.) (Figure 4A, right). The data show the mAb-SYT2 as a potential SVRM shuttle can transport functional molecules into neurons.

**Figure 4.**
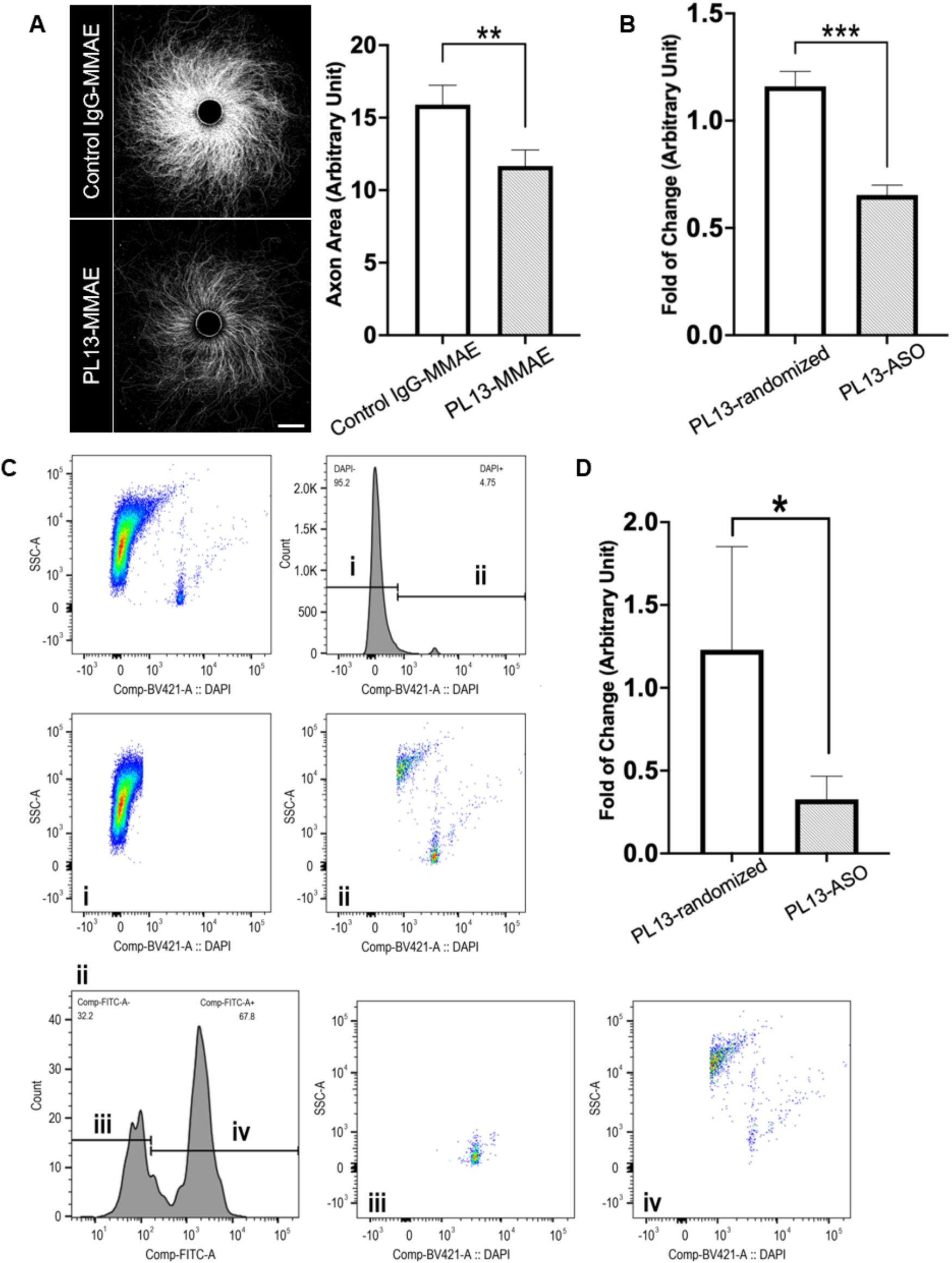
Delivery of an active payload to MN with SVRM antibody shuttle (A) Left shows a representation of PL13-MMAE delivery resulted in axon degeneration observed in the *in vitro* model. Scale bar 500µm. Right shows the graphical representation of PL13-MMAE delivery in the *in vitro* model showed 26.6% decrease in axon area compared to control chimera IgG-MMAE. (B) *in vitro* delivery of PL13-Malat1-ASO resulted in 34% knockdown of Malat1 RNA compared to delivery of PL13-randomized-ASO. (C) Mice were intravenously injected with PL13 conjugated with Malat1-ASO or randomized-ASO. Cells were gated for weak to strong DAPI staining (ii). DAPI positive cells (ii) were further sorted for PL13-positive (iv) or negative (iii) population. (D) PL13-positive cells from PL13-Malat1-ASO injected mice had 73.4% RNA knockdown compared to PL13-positive cells from PL13-randomized-ASO injected mice. Data expressed mean±s.e.m, n=3 and analyzed with t-test; \*\*\**p*<0.01, \*\**p*<0.05, \**p*≤0.1.

In addition to MMAE, the delivery of ASO to neurons was examined. We constructed *Malat1* gapmer ASO (Malat1-ASO) with DBCO-PEG4-Val-Cit-PAB-PNP linker. This linker construction with Malat1-ASO neither inhibited ASO activity (Figure S4A) nor did conjugation of PL13 with linker-ASO obstruct mAb-SYT2 binding efficiency to SYT2 antigen (Figure S4B). The conjugation rate of PL13 with linker-ASO was approximately 80% based on band intensity analysis with FIJI software (Figure S4C). Utilizing the same *in vitro* induced-presynapse model, PL13 conjugated with 0.6µM Malat1-ASO had a 34% decrease (0.6533±0.0464 s.e.m.) in Malat1 RNA expression compared to control PL13-randomized-ASO (1.160±0.0697 s.e.m.) (Figure 4B). These results showed that mAb-SYT2 transported functional small molecules and oligonucleotides into neurons.

Additionally, we sought to determine whether mAb-SYT2 was able to deliver functional ASO *in vivo*. Mice were intravenously injected with PL13-Malat1-ASO or PL13-randomized-ASO and the spinal cords were collected on day 10 post-IV injection for Malat1 RNA quantification. Because MNs are such a small population in the spinal cord, we isolated PL13-labeled cells from the spinal cord through cell sorting, which was achieved by additional PL13 IV administration at day 7 prior to sample collection at day 10. Spinal cord cells stained for DAPI were gated with FACS. PL13-positive cells were sorted by fluorescence staining of human Fc region of PL13 (Figure 4C). PL13-positive cells isolated from mice injected with PL13-Malat1-ASO had a notably Malat1 RNA reduction (0.327±0.1398 s.e.m.) compared to PL13-positive cells of mice injected with PL13-randomized-ASO (1.23±0.6228 s.e.m.) (Figure 4D). Similarly, PL13-positive cells showed a decrease in Malat1 RNA expression compared to PL13-negative cells isolated from the spinal cord of mice injected with PL13-Malat1-ASO (Figure S4D). These results indicate that PL13 delivery of Malat1-ASO was able to escape from the endo-lysosomal system into the cytoplasm and enter the nucleus for successful knockdown. Together, the *in vitro* and *in vivo* data signify that SVRM shuttle is capable of delivering therapeutic molecules to neurons. Specifically, when SYT2 is used as an antigen and administered intravenously, mAb-Syt2 can deliver therapeutic molecules through the MNs to the spinal cord.

## Discussion

In this study, we demonstrated the efficacy of antibodies as molecular shuttles that target the luminal domain of SV transmembrane protein enables neuronal delivery of small molecules and oligonucleotides through intrinsic SVRM. We employed spinal MN as a model system, capitalizing on their axon terminals are exempted from the BBB and BSCB regulation. Also this model has *in vivo* spatial separation of their somata located inside of these barriers and the synaptic terminals on the outside of these barriers. Our focus centered on SYT2, a protein abundantly expressed in skeletal muscle-projecting neurons, whose single-pass membrane structure renders it ideal for antibody generation.^29, 30^

Our findings reveal that systemic administration of mAb-SYT2 results in SVRM-mediated uptake at the axon terminal, analogous with botulinum neurotoxin (BoNT) transport mechanisms.^31^ In the spinal cord, mAb-SYT2 signals accumulate in ChAT-positive MNs, suggesting retrograde transport of antibody-containing SVs via endosomal and/or autophagosomal pathways, as evidenced by colocalization with the lysosomal marker LAMP1 in the soma.

Extending our observations to cranial MNs, we detected mAb-SYT2 signals in ChAT-positive brainstem nuclei, including the oculomotor, abducens, and facial nuclei. Interestingly, majority of intrinsic SYT2 expression, visualized with a polyclonal antibody, did not colocalize with mAb-SYT2 signals suggesting possible SYT2 competitive binding. However, the data more likely indicate differential recognition of intrinsic axonal SYT2 versus terminal-transported SYT2 localization.

Our data also showed that mAb-SYT2 utilizing SVRM was capable of delivering functional molecules into MNs and retrograde to the spinal cord. Miyashita SI *et al*. reported the usage of deactivated BoNT for peptide delivery into MNs, but there are still risks of residual toxicity from unknown toxin receptor molecules and production of neutralizing antibodies due to its immunogenicity.^32, 33^ Instead, the use of antibodies for therapeutic molecule delivery can circumvent these issues as antibodies are well-characterized molecules, host-compatible, and their clinical safety profiles are well-established in various conditions such as long-term multiple administrations.^34–36^

A critical challenge in neurodegenerative disease treatment and its development is to efficiently transport therapeutic molecules into the cerebrospinal tissue, where molecular permeability is tightly regulated by the BBB and BSCB. In recent years, transferrin receptor-mediated transcytosis and intrathecal injection have attracted attention to address these challenges. However, both approaches lack cell specificity once insde the BBB and BSCB.^37–39^ This potentially result in suboptimal therapeutic concentrations in target cells and raises the risk of adverse reactions in non-targeted cells.^40^

Additionally, intrathecal injection is invasive and burdensome for patients.^41^ In contrast, systemic administration of SV lumen-selective antibodies, e.g. mAb-SYT2, could overcome these limitations to target MNs in the spinal cord and brainstem. Thus, IV injection of mAb-SYT2 could serve as a molecular shuttle to spinal and brainstem MNs.

The principle underlying our approach extends to other SV transmembrane proteins. For example, Synaptophysin, the most abundant SV protein, presents an opportunity for potentially enhanced molecule targeting and delivery to spinal MNs. Furthermore, the administration of SV lumen-selective antibodies by intrathecal injections or engineering bispecific antibodies targeting transferrin receptors might allow specific targeting to distinct neurons based on SV proteins expression profile in targeted neurons; post-BBB entry.

In conclusion, antibody shuttle targeting SYT2 luminal domain demonstrates the feasibility of employing SV lumen-selective antibodies as neuron molecular shuttles via SVRM. By combining the selection of antibodies with diverse features and specific for luminal domains of SV transmembrane proteins expressed in different neuronal subtypes, we can achieve precise targeting of neurons while minimizing off-target effects. This can pave the way for new possibilities for developing tailored therapeutic molecule delivery systems as well as create targeted interventions for specific neurons, which allows for more effective and personalized treatments of neurological disorders. Future research in this direction could lead to a new generation of neuron-specific drug delivery platforms, potentially revolutionizing the field of neuropharmacology and improving patient outcomes.

### Limitations of the study

SYT2 expression is not confined to spinal MNs; it is also prominent in the brainstem, as corroborated by our immunostaining data and literature reports.^42^ Antibodies can migrate into cerebrospinal tissues following intravenous administration, indicating that both the target antigen and the administration method significantly influence cell selectivity.^43^ The mechanism of antibody transcytosis into CNS cells post-NMJ retrogradation requires further investigation to elucidate therapeutic molecule distribution when using mAb-SYT2 as a shuttle. Our data confirm mAb-SYT2 presence in endo-lysosomes, which might imply cellular stress or physiological inhibition risks, although no immediate adverse effects were observed in *in vivo* studies. Comprehensive risk management remains crucial during drug development, necessitating detailed studies to ensure safety and efficacy.

---

**Figure S1.**
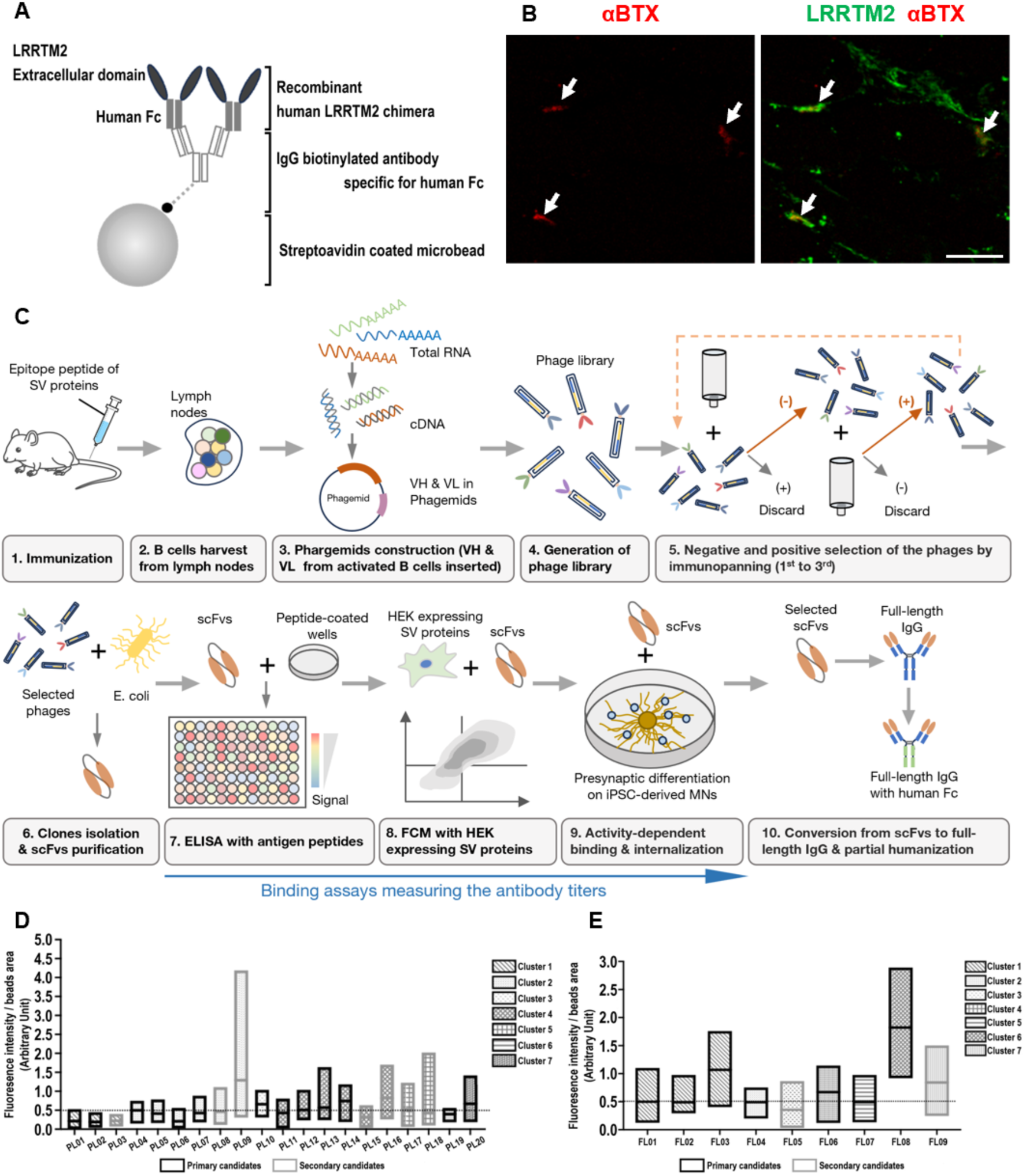
SVRM for molecular shuttling (A) Diagrammatic representation of presynapse induction microbeads coated with human LRRTM2 protein. (B) Localization of LRRTM2 at the neuromuscular junction in human skeletal muscle. (i) α-Bungarotoxin. (ii) Co-localization of LRRTM2 (labeled in green) with α-Bungarotoxin (labeled in red), arrow indicates NMJ. Scale bar represents 50µm. (C) Illustration for SVRM antibody shuttle screening, identification and generation of monoclonal antibodies specific for human and mouse SV luminal domain proteins. (D) Identification of scFVs with binding to SV proteins and internalization into SV during SVRM stimulation with 4-AP. Seven clusters from initial screening with FACS and ELISA were isolated. Graphs shows minimum and maximum value with mean value line. The median value of all the means (0.5) was drawn as a dotted line across the graphs. Further elimination of scFVs below the median of the mean values were done. scFV against full length SYT2 from AA1-62. FL03 and FL08, which has the highest internalization index to presynapse in the *in vitro* model were selected. Data were expressed in mean with minimum and maximum value, n>4. (E) scFV against partial length SYT2 from AA1-25. Clones with high, moderate and low internalization into presynapse in the *in vitro* model were selected. PL09 (high), PL10, PL20 (moderate) and PL13 (low) were selected for further study.

**Figure S2.**
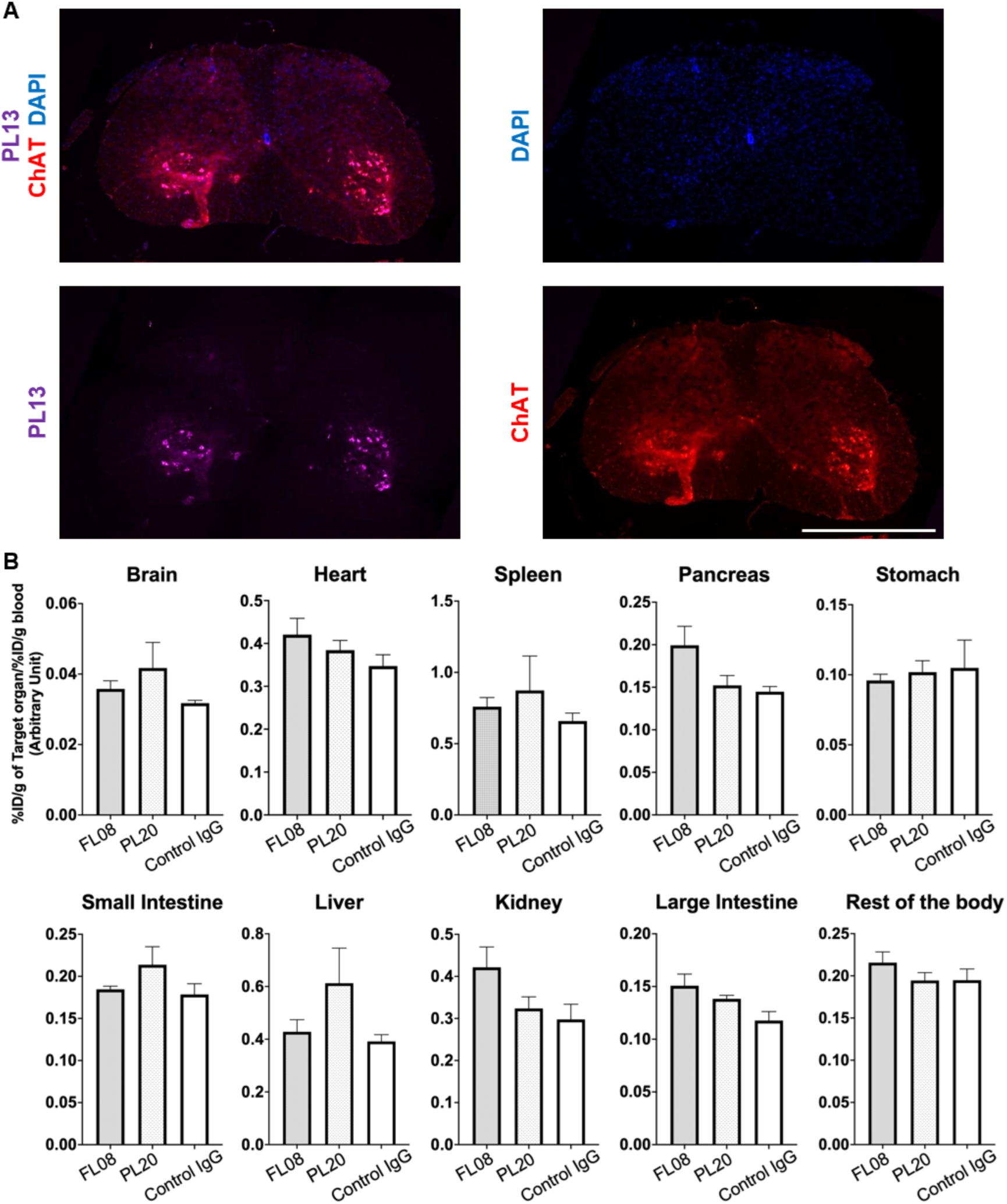
Intraspinal distribution of SVRM molecular shuttles after intravenous administration and RI results for other organs (A) PL13 (magenta) uptake into spinal cord after 72h. Scale bar represents 500µm (B) Localization ratio of FL08, PL20 and control IgG conjugated with radioisotope Zr^89^ in organs post intravenous injection.

**Figure S3.**
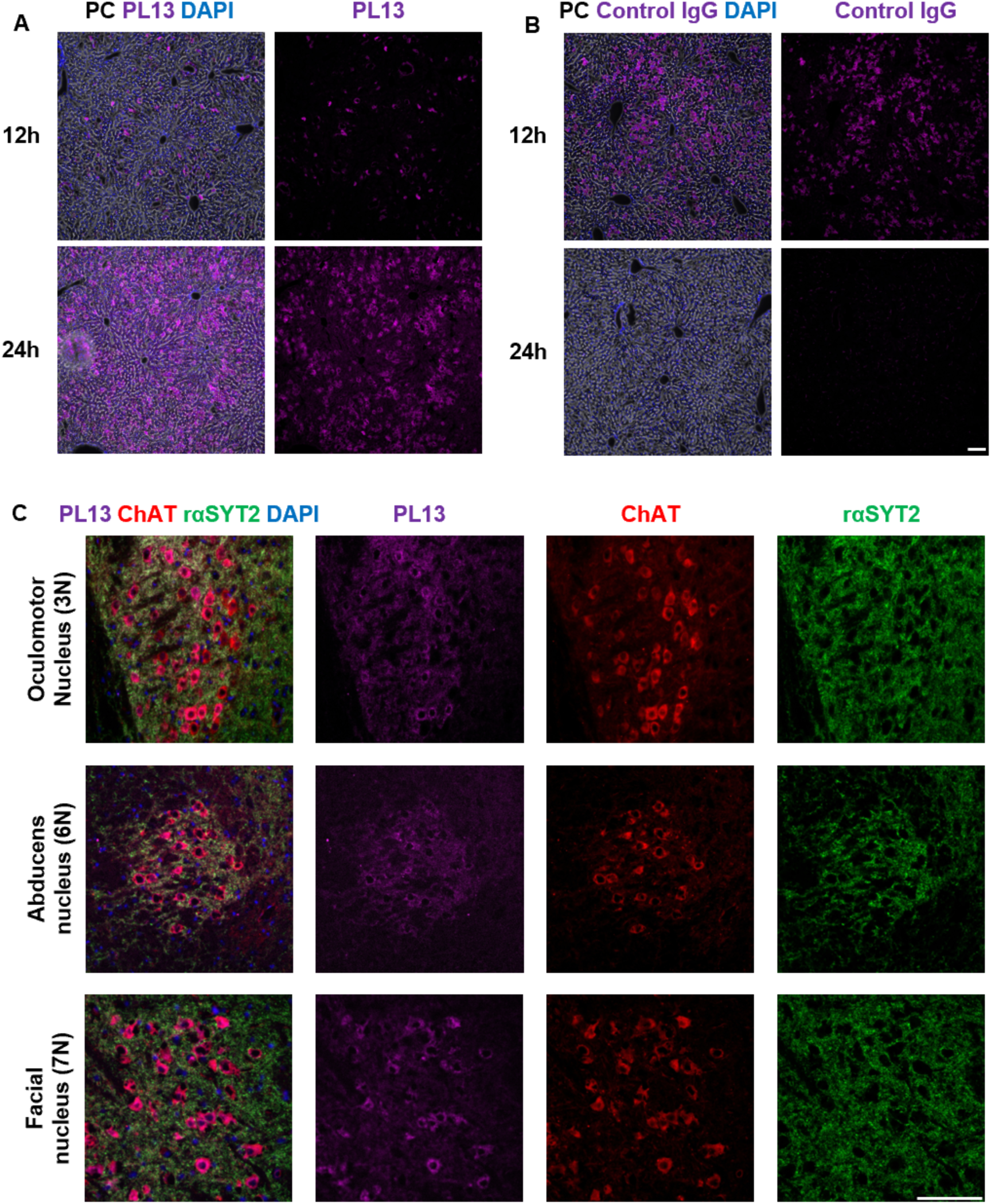
SVRM antibody shuttles distribution in liver and brainstem after intravenous administration Uptake of (A) PL13 (B) control IgG into liver and were detected up to 24h. Liver structures are viewed with phase contrast, antibodies in magenta and nucleus with DAPI. Scale bar represents 100µm. (C) Localization of PL13 (magenta) to brainstem 240h post-intravenous injection had co-localization with ChAT stained (red) MNs at 3N, 6N and 7N. Scale bar represents 100µm

**Figure S4.**
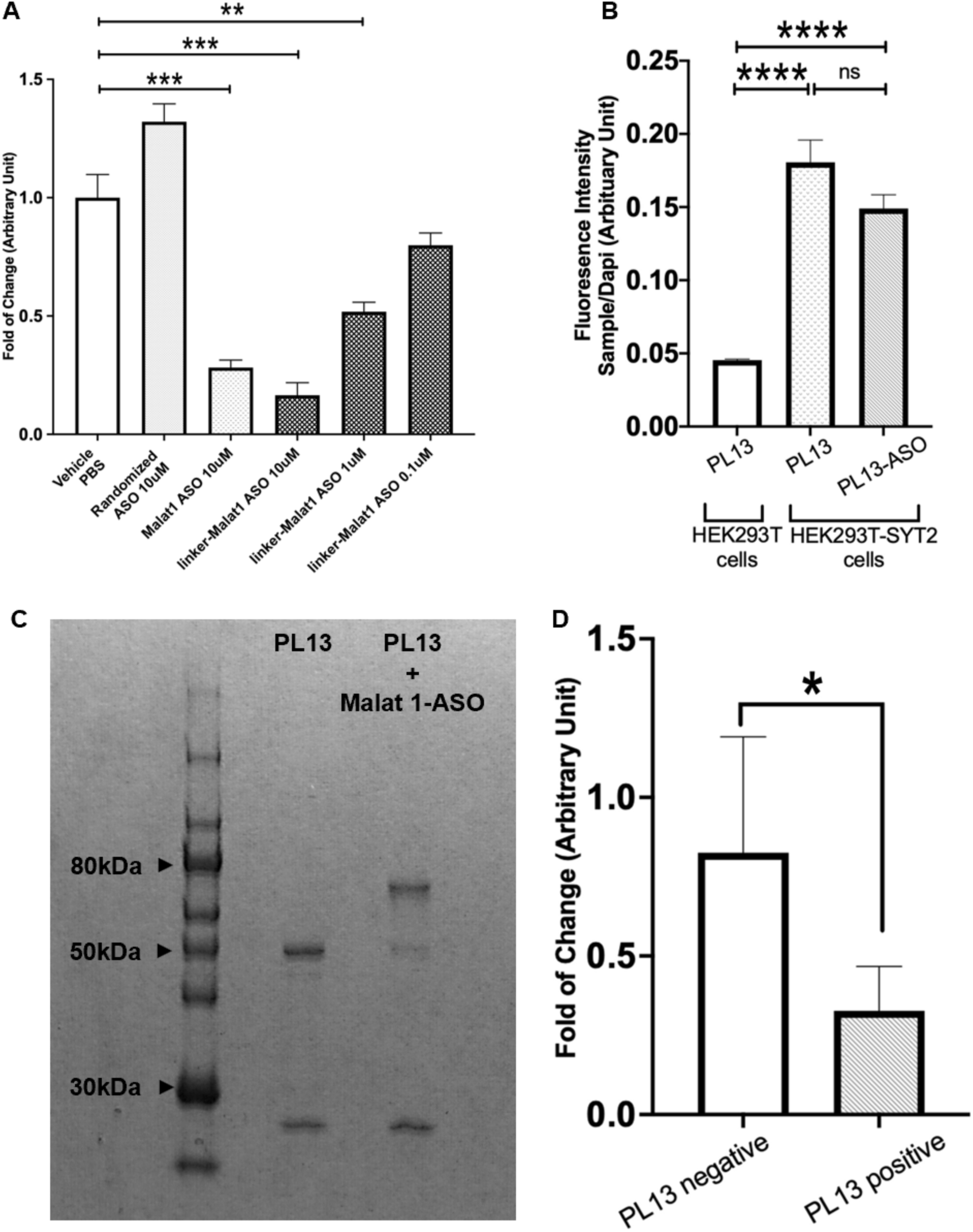
Characteristics and effects of PL13-Malat1-ASO (A) Evaluation of Malat1 ASO activity with and without linker attachment. Linker modified Malat1 ASO showed similar knockdown capabilities as Malat1 ASO alone. (B) Evaluation of PL13-ASO did not show obstruction of SYT2 luminal domain binding in HEK cells with SYT2 stable expression at the cell membrane. (C) PL13 conjugation to Malat1 ASO payload (approximately 70kDa) showed approximately 80% conjugation efficiency from band intensity measurements with FIJI. (D) Comparison of Malat1 RNA expression from cells isolated from spinal cord of mice injected with PL13-ASO. PL13 positive cells showed 60.4% decrease in Malat1 RNA expression compared to PL13 negative cells. Graphical data are expressed in mean±s.e.m., n=3 and was T-test analyzed; \*\*\*\**p*<0.001, \*\*\**p*<0.01, \*\**p*<0.05, \**p*≤0.1

**Table S1.**
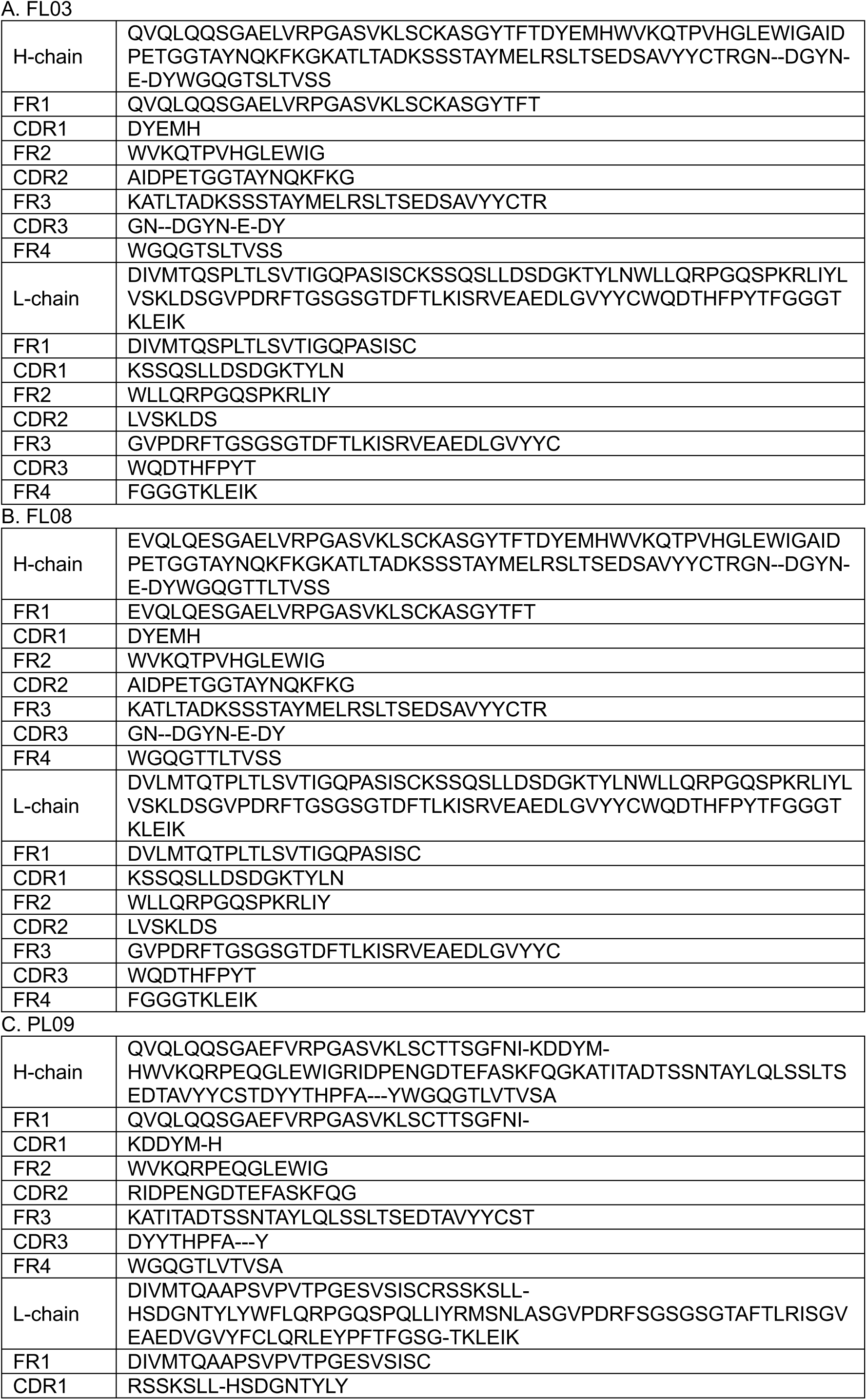
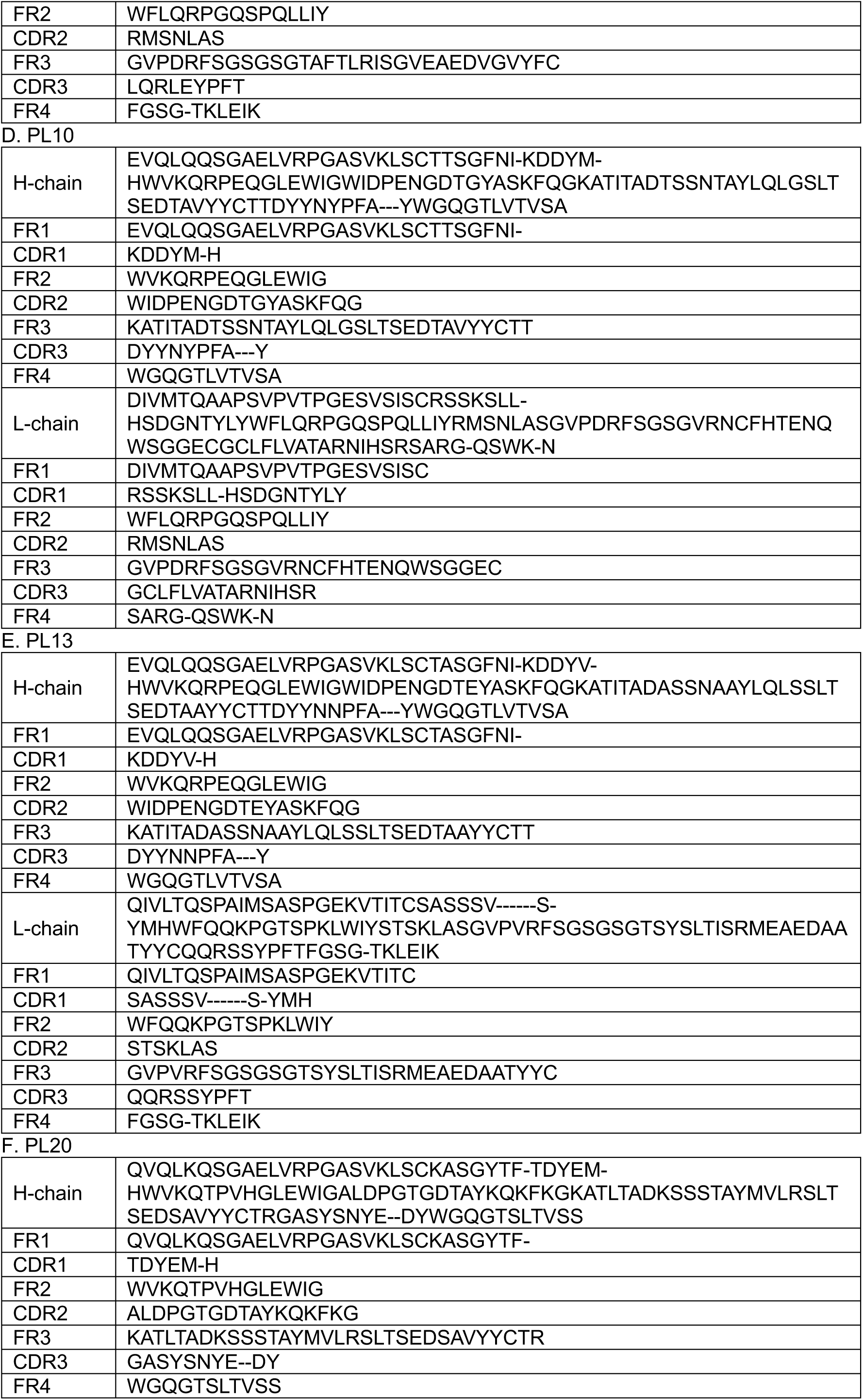
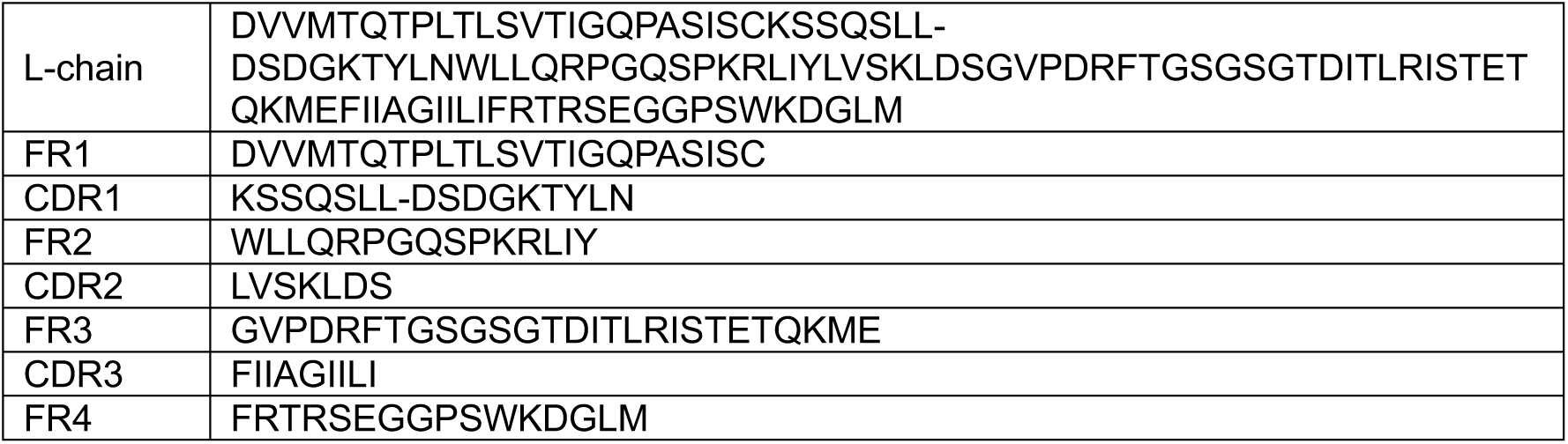
Sequences of FL and PL selected scFV clones.

**Table S2.**
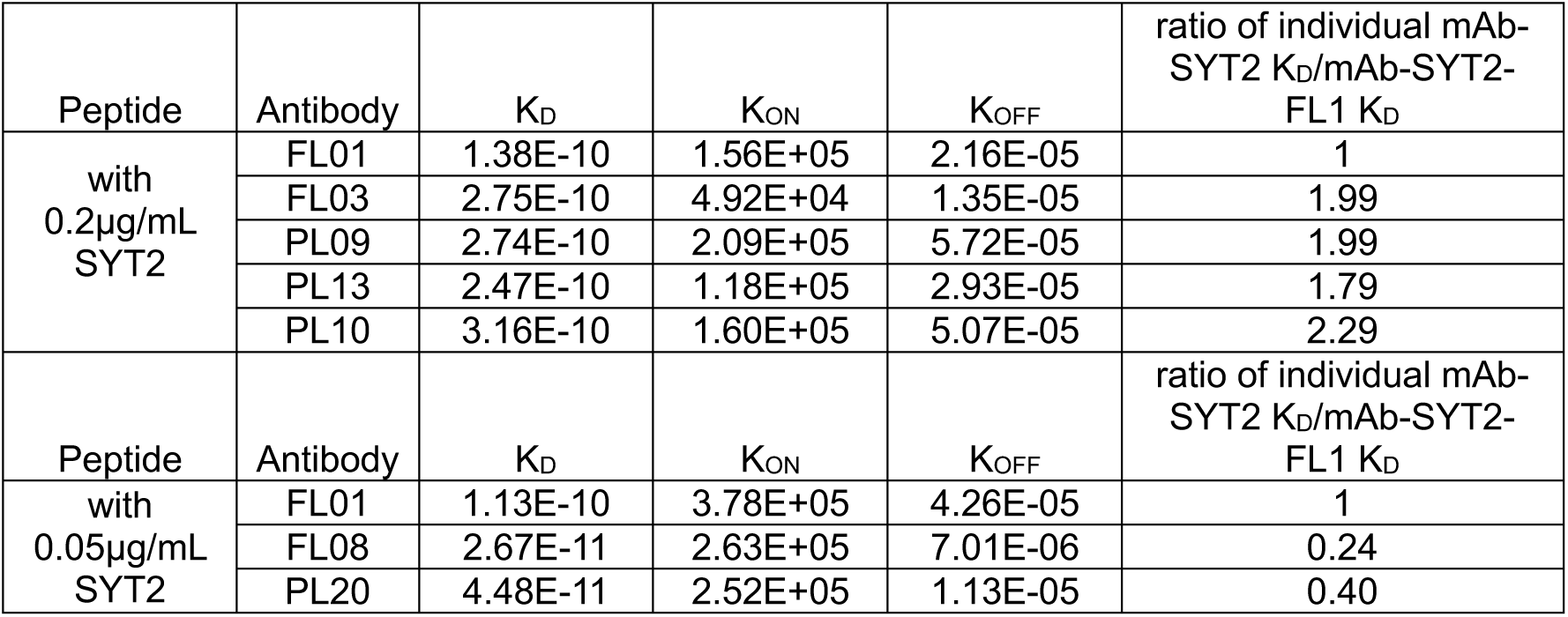
Dissociation constant and affinity characteristics of selected scFV that were constructed into chimeric full IgG with human Fc Constructed chimera antibodies were evaluated for their K_D_, K_ON_ and K_OFF_ properties. PL13 was characterized with moderate K_D_ ratio and as a SVRM molecular shuttle this characteristic will allow sufficient SYT2 binding and its release into the MN.

**Table S3.**
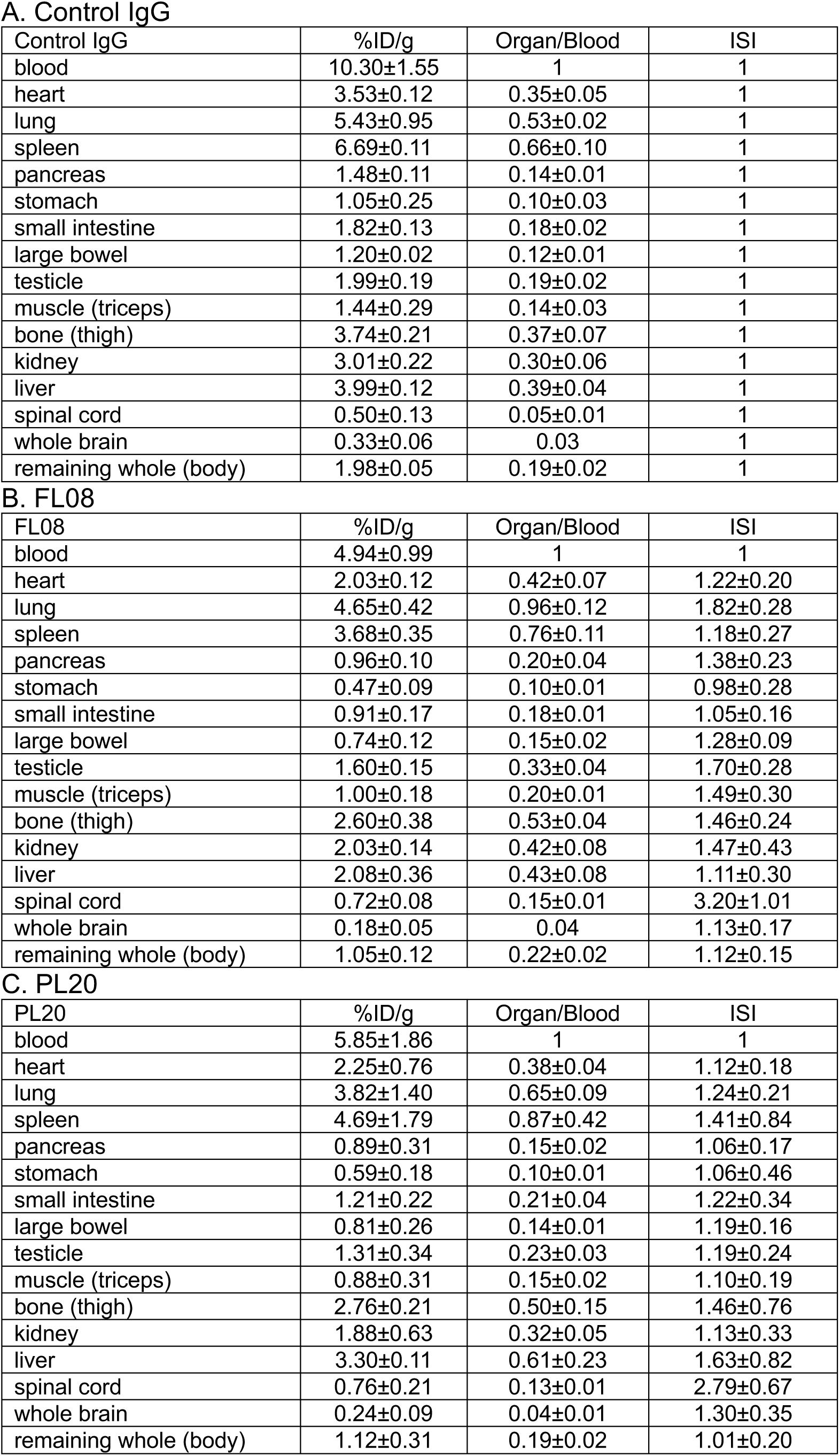
The localization ratio and immunospecificity index of mAb-SYT2 conjugated with Zr^89^. Spinal cord showed the highest immunospecificity index for FL08 and PL20. This indicates that mAb-SYT2 is specific and is efficient for spinal cord targeting.

A. Control IgG
| Control IgG | %ID/g | Organ/Blood | ISI |
| --- | --- | --- | --- |
| blood | 10.30±1.55 | 1 | 1 |
| heart | 3.53±0.12 | 0.35±0.05 | 1 |
| lung | 5.43±0.95 | 0.53±0.02 | 1 |
| spleen | 6.69±0.11 | 0.66±0.10 | 1 |
| pancreas | 1.48±0.11 | 0.14±0.01 | 1 |
| stomach | 1.05±0.25 | 0.10±0.03 | 1 |
| small intestine | 1.82±0.13 | 0.18±0.02 | 1 |
| large bowel | 1.20±0.02 | 0.12±0.01 | 1 |
| testicle | 1.99±0.19 | 0.19±0.02 | 1 |
| muscle (triceps) | 1.44±0.29 | 0.14±0.03 | 1 |
| bone (thigh) | 3.74±0.21 | 0.37±0.07 | 1 |
| kidney | 3.01±0.22 | 0.30±0.06 | 1 |
| liver | 3.99±0.12 | 0.39±0.04 | 1 |
| spinal cord | 0.50±0.13 | 0.05±0.01 | 1 |
| whole brain | 0.33±0.06 | 0.03 | 1 |
| remaining whole (body) | 1.98±0.05 | 0.19±0.02 | 1 |

B. FL08
| FL08 | %ID/g | Organ/Blood | ISI |
| --- | --- | --- | --- |
| blood | 4.94±0.99 | 1 | 1 |
| heart | 2.03±0.12 | 0.42±0.07 | 1.22±0.20 |
| lung | 4.65±0.42 | 0.96±0.12 | 1.82±0.28 |
| spleen | 3.68±0.35 | 0.76±0.11 | 1.18±0.27 |
| pancreas | 0.96±0.10 | 0.20±0.04 | 1.38±0.23 |
| stomach | 0.47±0.09 | 0.10±0.01 | 0.98±0.28 |
| small intestine | 0.91±0.17 | 0.18±0.01 | 1.05±0.16 |
| large bowel | 0.74±0.12 | 0.15±0.02 | 1.28±0.09 |
| testicle | 1.60±0.15 | 0.33±0.04 | 1.70±0.28 |
| muscle (triceps) | 1.00±0.18 | 0.20±0.01 | 1.49±0.30 |
| bone (thigh) | 2.60±0.38 | 0.53±0.04 | 1.46±0.24 |
| kidney | 2.03±0.14 | 0.42±0.08 | 1.47±0.43 |
| liver | 2.08±0.36 | 0.43±0.08 | 1.11±0.30 |
| spinal cord | 0.72±0.08 | 0.15±0.01 | 3.20±1.01 |
| whole brain | 0.18±0.05 | 0.04 | 1.13±0.17 |
| remaining whole (body) | 1.05±0.12 | 0.22±0.02 | 1.12±0.15 |

Table S3. The localization ratio and immunospecificity index of mAb-SYT2 conjugated with Zr<sup>89</sup>.
| PL20 | %ID/g | Organ/Blood | ISI |
| --- | --- | --- | --- |
| blood | 5.85±1.86 | 1 | 1 |
| heart | 2.25±0.76 | 0.38±0.04 | 1.12±0.18 |
| lung | 3.82±1.40 | 0.65±0.09 | 1.24±0.21 |
| spleen | 4.69±1.79 | 0.87±0.42 | 1.41±0.84 |
| pancreas | 0.89±0.31 | 0.15±0.02 | 1.06±0.17 |
| stomach | 0.59±0.18 | 0.10±0.01 | 1.06±0.46 |
| small intestine | 1.21±0.22 | 0.21±0.04 | 1.22±0.34 |
| large bowel | 0.81±0.26 | 0.14±0.01 | 1.19±0.16 |
| testicle | 1.31±0.34 | 0.23±0.03 | 1.19±0.24 |
| muscle (triceps) | 0.88±0.31 | 0.15±0.02 | 1.10±0.19 |
| bone (thigh) | 2.76±0.21 | 0.50±0.15 | 1.46±0.76 |
| kidney | 1.88±0.63 | 0.32±0.05 | 1.13±0.33 |
| liver | 3.30±0.11 | 0.61±0.23 | 1.63±0.82 |
| spinal cord | 0.76±0.21 | 0.13±0.01 | 2.79±0.67 |
| whole brain | 0.24±0.09 | 0.04±0.01 | 1.30±0.35 |
| remaining whole (body) | 1.12±0.31 | 0.19±0.02 | 1.01±0.20 |

## Materials and Methods

### Animals

For antibody distribution and ASO studies, we used 5 to 8-week-old female C57Bl/6 mice, and for RI studies, we used 5- to 7-week-old male ICR mice. All mice were purchased from Jackson Laboratory Japan Co., Ltd. Mice were housed in a temperature (22-24 °C) and humidity (40-51%) controlled room. Food and water were provided ad libitum, and the mice were housed under a 12-hour light/dark cycle. All behavioral manipulations were performed during the light phase. We complied with local and national ethical and legal regulations regarding the use of mice in research. All experimental protocols were approved by the animal testing regulations of KAC Co., Ltd. and Nihon Medi-Physics Co., Ltd., the animal experiment committee regulations, and the animal experiment approval regulations.

### Human Samples

The human study protocol was approved by the Tohoku University Hospital’s Institutional Review Board (Approval nos. 2022-1-144, 2022-1-848).

### Cells lines

HEK293T cells (CRL-3216, Lot:70041165) were purchased ATCC Inc. iPSC-derived Normal Human Motor Neurons (Lot:200238, 400329, 400445) were purchased iXcells Inc. Cultures were performed as described in METHOD DETAILS below.

### Preparation of Synaptic microbeads

The preparation of Synaptic microbeads was based on Yumoto et. al., (19). Briefly, SuperAvidin™ Coated Microspheres (Beads, Bangs Laboratories) were aliquoted at a concentration of 200 µg/mL and then washed with wash buffer (0.02% Albumin, Bovine Serum (BSA, NACALAI TESQUE) in PBS (WAKO)). Anti-Human IgG (Fc specific)-Biotin antibody, mouse monoclonal (Sigma Aldrich Technologies) at a concentration of 111 µg/mL was added, and then the microbeads were inverted and mixed at 4°C for 3 hours and then washed with wash buffer. Synaptic microbeads were added with Recombinant Human LRRTM2 Fc Chimera Protein, CF (R&D Systems) at a concentration of 66.6 µg/mL. Control microbeads were added with Native Human IgG FC fragment protein (Abcam) at a concentration of 66.6 µg/mL. Microbeads were mixed and inverted at 4°C for 3 hours and then washed with wash buffer. The final concentration of beads was 200 µg/mL.

### Cell culture (Motor neuron)

Human motor neurons (iXCells) were thawed and processed with Dead Cell Removal Kit (STEMCELL) according to manufacturer’s protocol. Cells were seeded onto a PrimeSurface® Plate96V (Sumitomo Bakelite) at 2×10^4^ cells/well. Then, the cells were incubated at 37℃ and 5% CO_2_ for one week to produce spheroids. During the cell culture, the medium was replaced twice using the Motor Neuron Culture Medium Kit (STEMCELL). EZVIEW Glass Bottom Culture Plates LB (IWAKI) were coated with 10 µg/mL poly-D-lysine hydrobromide (Sigma Aldrich Technologies) for one day (4°C), and then coated with 175 µg/mL Geltrex™ LDEV-Free, hESC-Qualified, Reduced Growth Factor Basement Membrane Matrix (Thermo Fisher Scientific) for three hours at 37°C with 5% CO_2_. Spheres were transferred from PrimeSurface® Plate 96V to EZVIEW Glass Bottom Culture Plates LB. The cells were cultured for one month, changing the medium three times a week. Cells were cultured in Neurobasal Plus Medium (Thermo Fisher Scientific), 2% B-27 Plus Supplement (50X) (Thermo Fisher Scientific), 1%Penicillin-Streptomycin (Thermo Fisher Scientific), 20 ng/mL Brain-derived neurotrophic factor (BDNF, GenScript), 20 ng/mL Glial Cell Line-derived Neurotrophic Factor (GDNF, GenScript) in a 5% CO_2_ atmosphere at 37 °C. After culturing for one month, beads were added and the cells were cultured for another 48 hours to promote synapse formation, after which they were used for testing.

### Cell culture (HEK293T)

We used the commercial HEK293T cell line (ATCC). SYT2 stable expression cell lines were generated at MBL. Cells were grown in DMEM (Gibco) supplemented with 1% Penicillin-Streptomycin (Thermo Fisher Scientific), 10% FBS (Gibco), 1% GlutaMAX™ I (Thermo Fisher Scientific) in a 5% CO_2_ atmosphere at 37 °C.

### Human tissues study

Thirty-micrometer-thick transverse cryosections were cut on a cryostat (Leica), collected onto silanized slides, and stored at −80°C. The sections were fixed with ice-cold acetone for 10 min, and dried. After washing in Tris-buffered saline (TBS), pH 7.4, nonspecific binding was blocked with 5% normal goat serum (Vector) and 0.3% Triton X-100 in TBS for 20 min at room temperature. The sections were then incubated with primary antibody (LRRTM2, Novus Biologicals) and alpha-Bungarotoxin, Alexa594 conjugate (Thermo Fisher Scientific) in the blocking solution for 24 h at 4°C and then rinsed extensively in TBS, followed by incubation with the appropriate secondary antibodies conjugated to fluorochromes (Alexa Fluor 488, Thermo Fisher Scientific) in Antibody Diluent (Dako) overnight at 4°C. Subsequently, the slides were washed extensively in TBS, dipped into distilled water, coverslipped with PermaFluor (Thermo Fisher Scientific), and stored in the dark at 4°C. We analyzed the fluorophore-labeled sections under a confocal laser scanning microscope system with Ar (488 nm), and HeNe-green (594 nm) laser units (IX 71 and FV300; Olympus, Tokyo, Japan). Digital images were captured using acquisition software (Fluoview version 4.3; Olympus).

### Rabbit SYT2 uptake study

Cells were maintained in Neurobasal Plus Medium (Thermo Fisher Scientific), 2% B-27 Plus Supplement (50X) (Thermo Fisher Scientific), 1%Penicillin-Streptomycin (Thermo Fisher Scientific), 20 ng/mL BDNF(GenScript), 20 ng/mL GDNF(GenScript). This study used 10 µg/mL synaptotagmin 2 (SYT2) antibody luminal domain (Synaptic systems), 10 µg/mL normal rabbit IgG control (R&D Systems), and 10 µg/mL synaptotagmin 2 antibody cytoplasmic domain (Synaptic systems). To enhance antibody uptake, a mixture of antibody and 100 µM 4-aminopyridine (Sigma-Aldrich) was added and incubated at 37°C for 30 minutes, then washed with culture medium. Finally, cells were fixed with 2% PFA (NACALAI TESQUE).

### Immunofluorescence staining (Cell)

Cells were fixed in 2% paraformaldehyde (PFA, NACALAI TESQUE) and then washed in wash buffer (0.02% Triton(R) X-100 (Tx, NACALAI TESQUE) in PBS (WAKO)). Cells were then incubated in permeabilizing buffer (0.25% Tx in PBS) and blocking solution (2% Normal Goat Serum (NGS, Thermo Fisher Scientific), 1%BSA (NACALAI TESQUE), 1% Fetal Bovine Serum (FBS, Gibco), 0.02% Tx in PBS) at room temperature for 1 h with gentle agitation. Detection of targets was performed by incubating cells in a cocktail of primary antibodies for a day at 4°C under agitation. The cell was then washed with wash buffer and incubated with Alexa Fluor-conjugated secondary antibodies (Invitrogen) for 1 h at room temperature under agitation. After additional rinses in wash buffer, Cells were treated with 1% PFA for re-crosslinking. Primary antibody cocktails were comprised of antibodies from different species or different isotypes of IgG. A list of primary antibodies used in this study and their respective dilutions can be found in KEY RESOURCES TABLE. Microscope (Leica DMi8), Camera (DFC9000GTC) and Filter (XLED-Q-P) were used for imaging. Image J v1.53e and Meta Morph version 7.10.2.240 were used for analysis.

### Antibodies generation

Antibody production was outsourced to MEDICAL & BIOLOGICAL LABORATORIES CO., LTD. (MBL). Briefly, the immunogen and panning antigens used for antigen preparation were synthesized by Beijing SciLight Biotechnology Ltd. The sequences of the synthesized peptides for immunogen and panning are as follows. <PEPTIDES for immunogen> h2F-Ag (1-62aa), MRNIFKRNQEPIVAPATTTATMPIGPVDNSTESGGAGESQEDMFAKLKEKLFNEINK IPLPP-C. h2A-Ag (1-25aa), MRNIFKRNQEPIVAPATTTATMPIG-C. <Peptides for panning> h2F-bio (1-62aa), MRNIFKRNQEPIVAPATTTATMPIGPVDNSTESGGAGESQEDMFAKLKEKLFNEINK IPLPP-K-biotin. m2F-bio (1-65aa), MRNIFKRNQEPNVAPATTTATMPLAPVAPADNSTESTGPGESQEDMFAKLKEKFFNEINKIPLPP-K-biotin. h1F-bio (1-58aa), MVSESHHEALAAPPVTTVATVLPSNATEPASPGEGKEDAFSKLKEKFMNELHKIPLPP-K-biotin.

KLH (Keyhole Limpet Hemocyanin) was conjugated to the antigen-conditioned peptide and the concentration was prepared to 1.0 mg/mL. The solution was immunized once a week for a total of four times to two each of Balb/c mice and diseased mice. Serum titers against the peptide antigen were evaluated by ELISA of the immunized mice sera. The reactivity of SYT2 Antibody luminal domain (Synaptic systems) to biotinylated-h2F peptide was also compared. Next, we established cells stably expressing SYT2 antigen. We used the entire sequence of human synaptotagmin 2 (419AA, https://www.uniprot.org/uniprotkb/Q8N9I0/entry) and mouse synaptotagmin 2 (419AA, https://www.uniprot.org/uniprotkb/P46097/ entry) were transfected with pCMV6-AC-Myc-DDK-IRES-GFP-Puro Mammalian Expression Vector (OriGene Technologies, Inc.) vector, respectively, and cultured for 3 days.

After GFP expression was confirmed by fluorescence microscopy, selection was performed in medium containing 2.5 µg/mL of puromycin. The selected cells were subjected to FCM analysis and confirm expression of SYT2 antigen. Cells without vector were used as a negative control for cell samples. Antibody phage libraries were then generated, and antibody selection was performed on h2F-KLH- and h2A-KLH-immunized mice, respectively, followed by panning. Antibody clones obtained by panning were induced with IPTG (isopropyl β-D-thiogalactopyranoside) and culture supernatants containing polyclonal scFv (single-chain variable fragment) were obtained. ELISA assay of polyclonal scFv was performed to evaluate the enrichment of peptide antigen-specific clones. h2F and h2A antibody clones obtained in the third panning operation were seeded onto plates for cloning. 96 colonies each from h2F and h2A libraries were cultured and IPTG-induced the culture supernatant containing monoclonal scFv was obtained by IPTG induction. ELISA assay was performed for monoclonal scFv. Antibody gene sequences of clones that tested positive by ELISA were analyzed by Azenta US, Inc. The amino acid sequence of scFV was confirmed from the results of gene analysis. Clones were clustered by amino acid sequence homology. Clones selected from ELISA evaluation and antibody gene analysis were IPTG induced and culture supernatant was concentrated 10-fold by ammonium sulfate precipitation. scFV FCM measurement was performed to evaluate its titer against hSYT2 antigens. scFV was subjected to ELISA and FCM evaluation and after IPTG induction, the culture supernatant was concentrated 10-fold by ammonium sulfate precipitation. The concentrated supernatant was dialyzed and filtered through a 0.22 um filter and used for the scFV uptake test. This test was performed as in Rabbit SYT2 uptake test. Finally, seven clones were purified to Human-Mouse Chimera IgG1 antibody. The antibody used in this study was FL08, PL13 and PL20.

### Kinetics Measurement

Kinetics Measurement was outsourced to MEDICAL & BIOLOGICAL LABORATORIES CO., LTD. (MBL). For the antibodies produced, affinity (K_D_), binding kinetics (K_ON_) and dissociation kinetics (K_OFF_) to biotinylated h2F peptides were measured. Instrument used for the measurements was an Octet RED96e system (SARTORIUS). The biosensor used was Octet streptavidin Biosesor (SARTORIUS). Concentrations of the measured samples were 5, 2.5, 1.25, 0.63, 0.31, 0.17, 0.08, and 0 nM. Sensitizing antigen was 0.2 µg/mL or 0.05 µg/mL. Buffer was 0.02% Tween-20/ 0.01% BSA/PBS.

### Antibody distribution (*in vivo*)

The animals used and the breeding conditions were described in the EXPERIMENTAL MODEL AND STUDY PARTICIPANT DETAILS. In this study, PL13 antibody or Human IgG1 isotype control chimeric mAb (MBL) was used, and i.v. injected at a dose of 5 mg/kg was done with a single injection. The mice were fixed, and organs were harvested at each time point (12h, 24h, 72h and 240h after i.v. injection). Under isoflurane inhalation anesthesia, the mice were perfused with PBS into the left ventricle and euthanized. After confirming that the blood in the body had been drained, PBS was replaced and fixed with 4% PFA. After perfusion, each organ was harvested from the mice, and blood, etc. were washed away with PBS, and then each organ was immersed in 4% PFA for one day. The tissue was then sucrose-substituted with 10% sucrose (WAKO) for 2 hours (4°C), 20% sucrose for 2 hours (4°C), and 30% sucrose overnight (4°C), and embedded in OCT compound (Sakura Finetek Japan). Sections were made with a Leica cryostat and were 10 µm thick.

### Immunofluorescence staining (tissue)

Sections were washed in wash buffer (0.1% Tx (NACALAI TESQUE) in PBS (WAKO)). Sections were incubated in permeabilizing buffer (0.25% Tx in PBS) and then blocking solution (3% NGS (Thermo Fisher Scientific) or 3% Normal donkey serum (NDS, Merck), 2%BSA (NACALAI TESQUE), 0.1% Tx in PBS) at room temperature for 1 h. Detection of targets was performed by incubating sections in a cocktail of primary antibodies for a day at 4°C under agitation. The tissue was then washed with wash buffer and incubated with Dylight or Alexa Fluor-conjugated secondary antibodies (Invitrogen) and or 1.43 µM of DAPI (Invitrogen) for 1 h at room temperature. After additional rinses in wash buffer, sections were mounted with Fluoromount-G (Southern Biotechnology Associates). Microscope: Leica DMi8, Camera: DFC9000GTC, Filter: XLED-Q-P was used for the imaging. Meta Morph version 7.10.2.240 and Image J v1.53e software were used for analysis.

### Radioisotope (RI)

The RI test was outsourced to Nihon Medi-Physics Co., Ltd. Briefly, the study involving the use of animals in this study was conducted in compliance with the relevant regulations and standards after the experimental plan was designed in accordance with the Regulations on Safety Management of Biological Experiments of the Research Facilities of Nihon Medi-Physics Co. A total of 12 mice with normal ICR were obtained from Jackson Laboratory Japan, including one spare mouse per group for a total of 3 mice per antibody (total of 9 mice per group) for this study. In this study, we evaluated radioisotope Zr^8^9 labeling of the prepared FL08, PL20 and Human IgG1 isotype control chimeric antibody (Control IgG, MBL) by PET imaging of Zr^89^-labeled FL08, PL20 and Control IgG. In the Zr^89^-labeling study, Zr^89^-labeled FL08, PL20 and Control IgG antibodies were administered intravenously to the mice. In the Zr^89^-labeling study, CCAP (Chemical Conjugation by Affinity Peptide) modified antibody solutions of the prepared FL08, PL20 and Control IgG and investigated 89Zr conjugation methods including Zr^89^-labeling and quality evaluation using CCAP-modified antibody solutions. Next, PET imaging evaluation was performed using normal mice treated with Zr^89^-labeled FL08, PL20 and Control IgG antibody over time.

### Monomethyl auristatin E (MMAE) Uptake study

Cell culture was performed as described in Cell culture (Motor neuron). The binding of VcMMAE (mc-vc-PAB-MMAE) (MedChemExpress) to each antibody was performed using MagicLink™ Protein Protein Crosslinking Kit (BroadPharm) as manufacturer’s protocol. The binding ratio of antibody to MMAE is 1:2. The MMAE uptake test was performed as in Rabbit SYT2 uptake test. The final concentration of each reagent was 10 µg/mL. Microscope (Leica DMi8), Camera (DFC9000GTC) and Filter (XLED-Q-P) were used for imaging. Image J v1.53e was used for analysis. Primary and secondary antibodies are listed in Supplement table.

### AOC *in vitro* study

#### Generation of Antibody Oligonucleotide Conjugates (AOC)

For AOC generation, nucleic acid synthesis with linkers was outsourced to Ajinomoto Bio-Pharma to synthesize anti malat1 ASO (C6-(5’-) GmCATTmCTAATAGmCAGmC (−3’)) and randomized ASO (C6-(5’-) TmCAmCTmCGAAmCAGTAGT (−3’)). ASO and DBCO-PEG4-Val-Cit-PAB-PNP (BroadPharm, #BP-24520) were then conjugated as per protocol using oYo-Link® Azide (AlphaThera) and LED PX2 Photo-Crosslinking Device (AlphaThera). The binding ratio of antibody to the synthesized nucleic acids with linkers was 1:3 (molar ratio). Binding was confirmed by SDS-PAGE.

#### AOC binding evaluation study

Cells were seeded at 1×10^4^/well on Geltrex™ hESC-Qualified, Ready-To-Use, Reduced Growth Factor Basement Membrane Matrix (Thermo fisher scientific)-coated Black Microplate Flat Bottom 96Well, I Type gamma Ray Sterilized (AS ONE). The test was performed when the cells were 100% confluent. Each reagent was dissolved in medium and prepared to a final concentration of 1 µg/mL was incubated at 37°C, 5% CO_2_ for 1 hour, then washed with medium. The cells were fixed at a final concentration of 2% PFA and then immunofluorescence stained. Fluorescence intensity was measured at GloMax® Discover Microplate Reader (Promega).

#### AOC RNA knockdown study

For HEK293T experiment, Cells were seeded at 4×10^5^/well on CELL CULTURE MULTIWELL PLATE, 6 WELL, PS, CLEAR (Greiner). The test was performed when the cells were Approximately 70% confluent. Randomized Antisense Oligonucleotides (ASO) and Malat-1 ASO were prepared to a final concentration of 10 µM. DBCO Linker conjugate Malat-1 ASO was prepared at 0.1 µM, 1 µM and 10 µM. The incubation conditions were 37°C, 5% CO_2_ for a duration of 2 days. The motor neuron experiment, cell culture was performed as described in Cell culture (Motor neuron). This study was performed as described in Rabbit SYT2 uptake test. Conjugated PL13-Malat1-ASO or Malat1-ASO alone were added to the cells at final concentrations of 10 µg/mL and 2 µg/mL and incubated at 37°C for 2 days. In each experiment, cells were washed with PBS (WAKO) after incubation and collected in 0.125% Trypsin Solution (NACALAI TESQUE). RNA extraction, cDNA synthesis, and qPCR are described in “Quantitative PCR for RNA expression.”

#### AOC *in vivo* study Animal study

This test was outsourced to KAC Co., Ltd. The animals used and the breeding conditions were described in the EXPERIMENTAL MODEL AND STUDY PARTICIPANT DETAILS. Briefly, this study was based on Beaudet MJ, et al. Sci Rep. 2015. In this study, C57Bl/6 mice purchased from Jackson River were used. Each group had 3 mice and a total of 18 mice (N=3). PL13-Malat1-ASO conjugated or PL13-randomized-ASO conjugated was used, and i.v. injected at a dose of 5 mg/kg respectively, and then for Fluorescence-activated cell sorting (FACS), antibody was administered i.v. at 5 mg/kg 72 hours before sampling. Spinal cord cells were isolated using the Papain Dissociation System (Worthington Biochemical Corporation) according to the protocol. The isolated spinal cord cells were fixed with 2% PFA.

#### FACS

FACS was based on David Martin et. al., (44). Isolated cells were washed with PBS (WAKO), and cell counts were performed using Countess 3 (Thermo Fisher Scientific). Antibody solutions were prepared by diluting Goat anti-Human IgG (H+L) Cross-Adsorbed Secondary Antibody, Alexa Fluor™ Plus 488 (20 µg/mL, Invitrogen) and DAPI (1.43 µM, Invitrogen) with staining buffer (1mM DTT (NACALAI TESQUE), 0.1U/µL RiboLock RNase Inhibitor (Thermo Fisher Scientific), 0.1% Tween (NACALAI TESQUE) in PBS(WAKO)). The cell concentration was adjusted with antibody solution to 1×10^5^/50 µL. The adjusted antibody solution was inverted and mixed at 4°C for 30 minutes in a light-shielded environment. Stained cells were washed with wash buffer (0.1% Tween in PBS). Finally, the cells were diluted with staining buffer to a cell concentration of 1×10^7^/mL and used. To identify Malat1 expression, stained spinal cord cell suspensions were sorted for PL13 and DAPI-positive cells using SORPAria or Aria 3 cell sorter (BD). The analysis software used was FlowJo, LLC (BD).

#### Quantitative PCR for RNA expression

For qPCR analysis, total RNA was extracted using the NucleoSpin® totalRNA FFPE XS (MACHEREY-NAGEL) or RNeasy Plus Mini Kit (Qiagen), and reverse-transcribed with the QuantAccuracy®, RT-RamDA® cDNA Synthesis Kit (TOYOBO). Gene expression was analyzed by qPCR with TB Green® Premix Ex Taq™ (Tli RNaseH Plus) (Takara Bio). The PCR primers used in this study are listed in the key resources table.

#### Quantification and statistical analysis

Analyses were performed using Prism 9.0 (GraphPad), and graphs were generated using Prism 9.0. Data represents mean ± SD. Specific tests (e.g., t-test, one-way ANOVA tests) and significance are indicated in figure legends.

#### Materials List

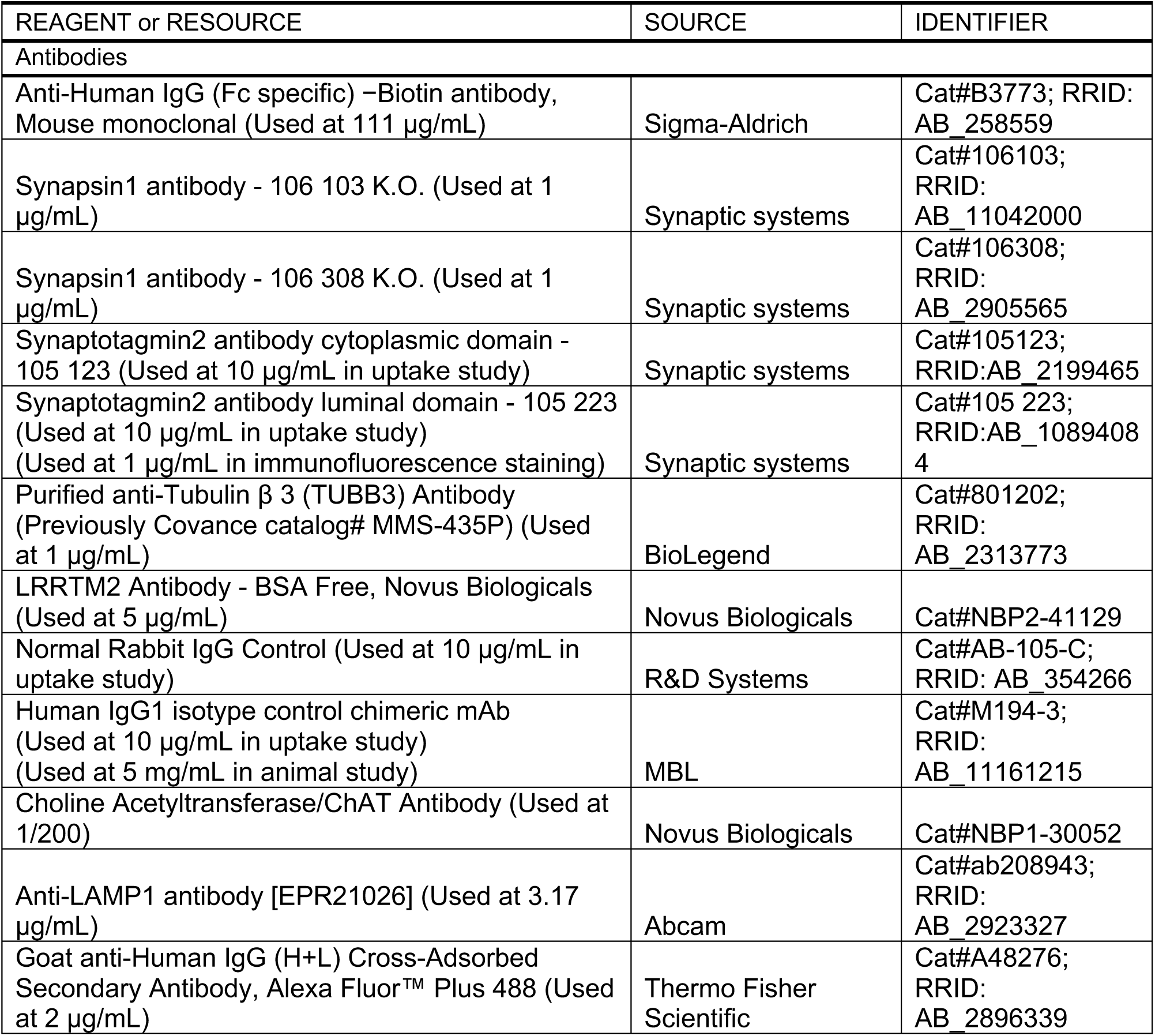
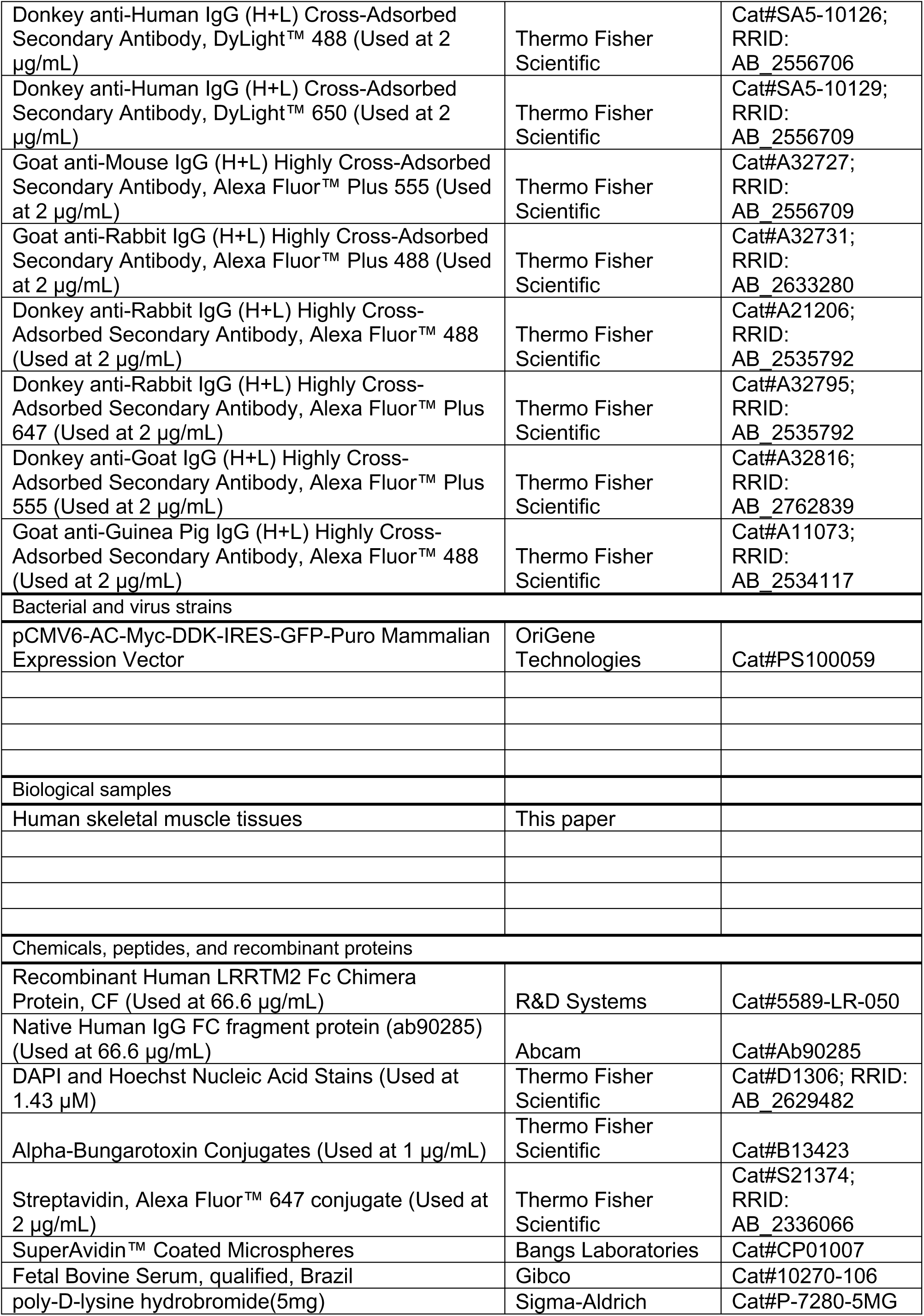
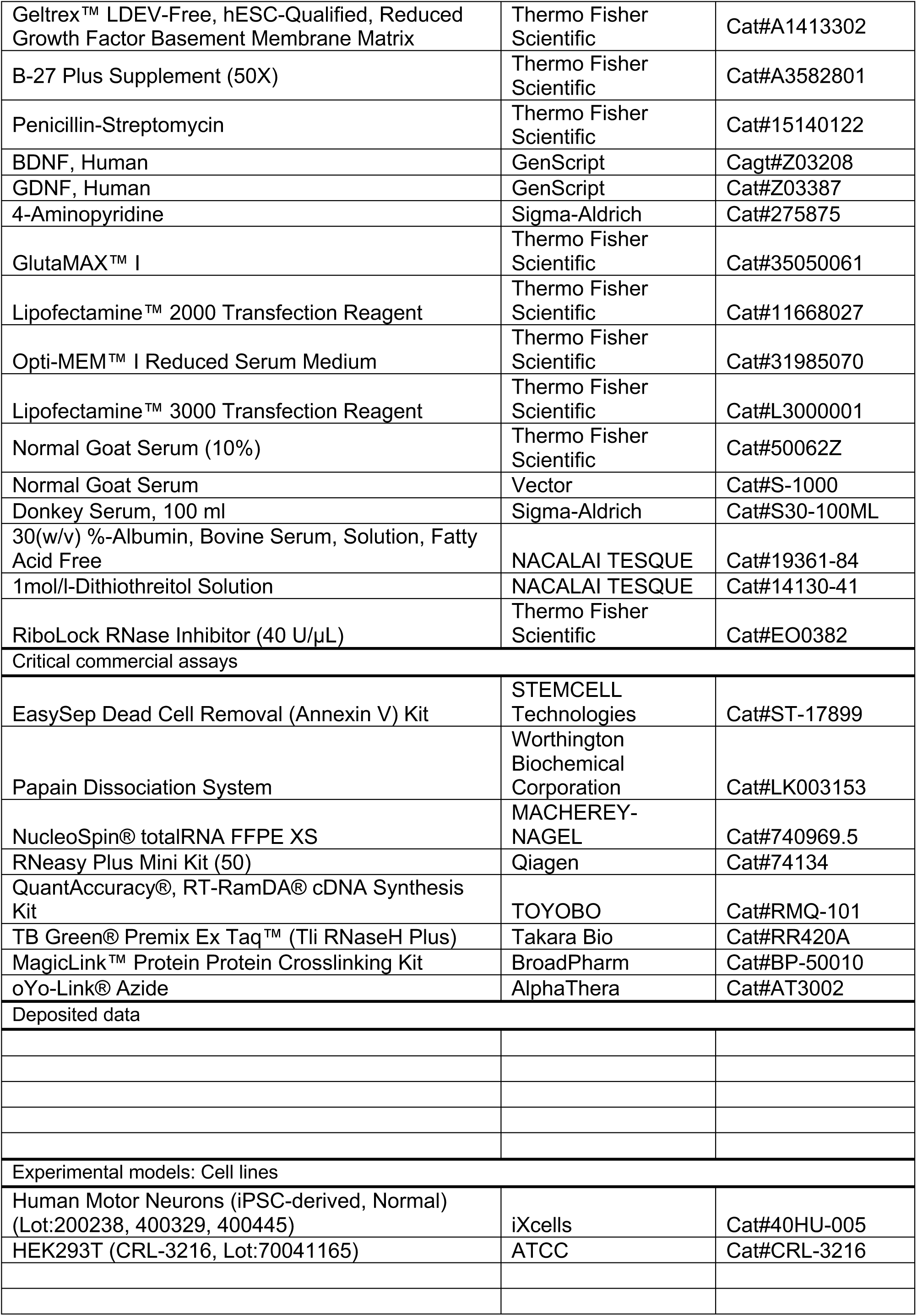
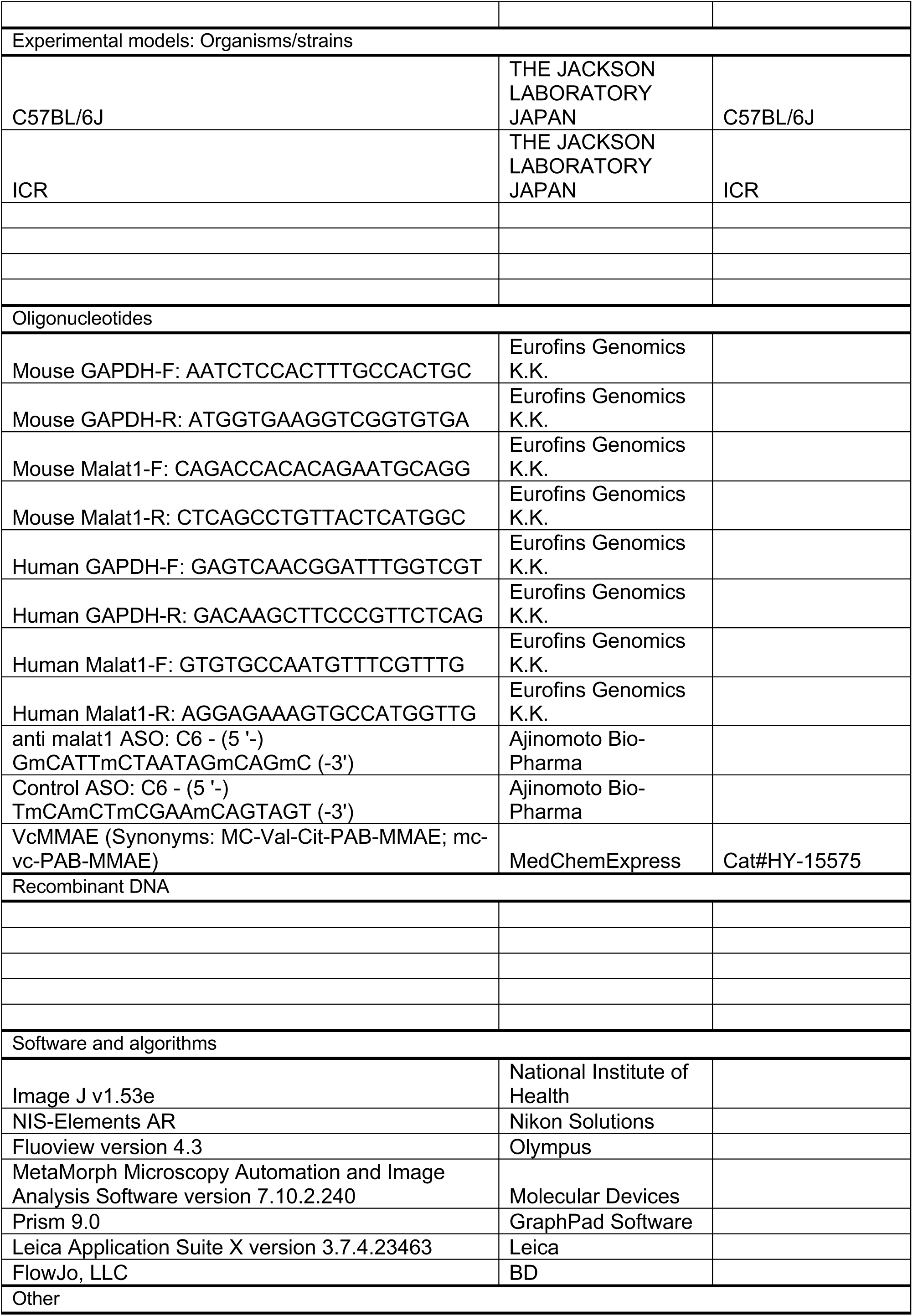
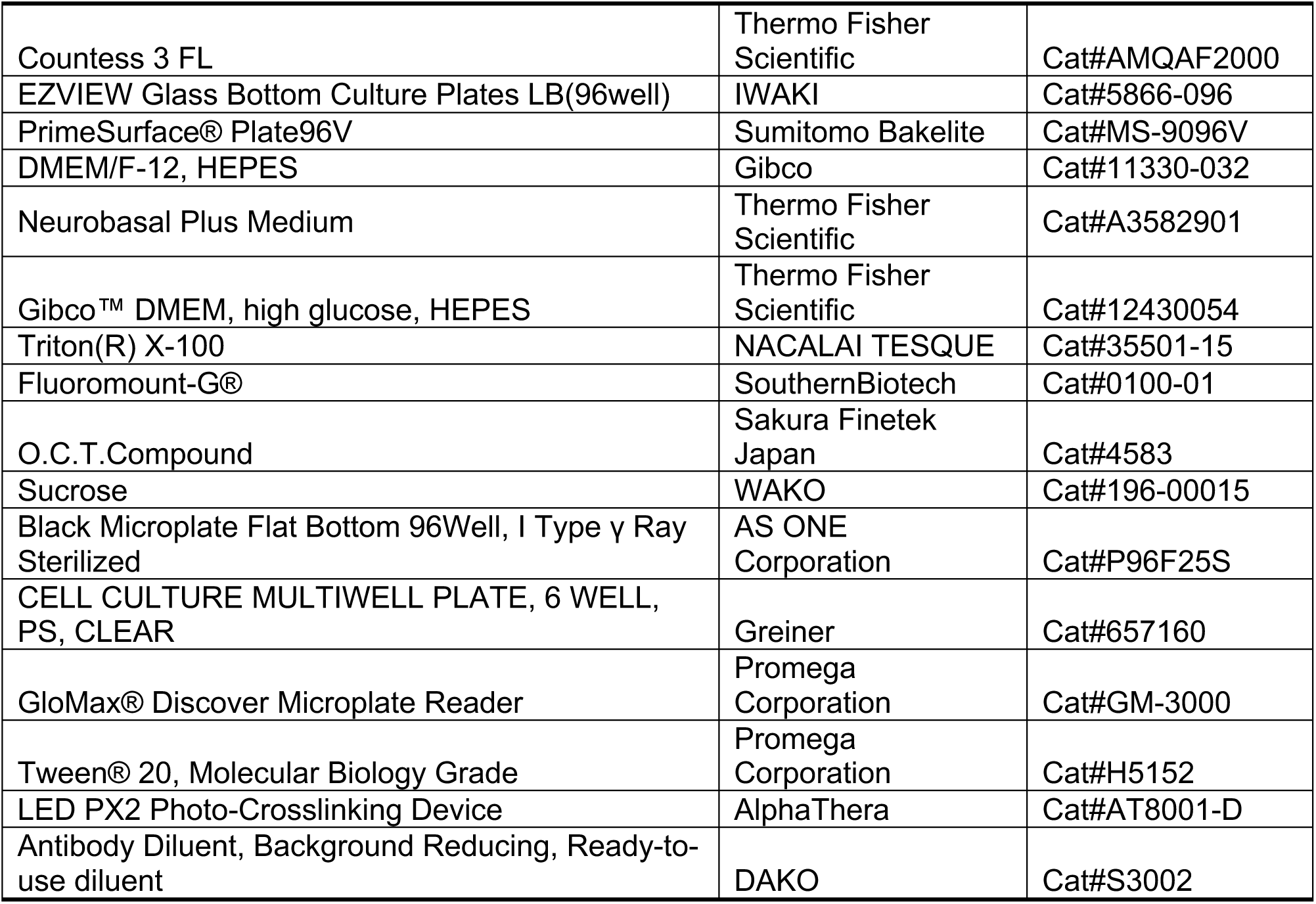

## ACKNOWLEDGMENTS

Flow cytometry was performed in the IMSUT FACS Core laboratory. We acknowledge the IMSUT FACS Core laboratory for assistance with flow cytometry analysis. We thank Mami Okamoto, Sanae Ishizuka, Tomomi Sakamoto, Rumiko Izumi, Masako Suzuki, Mai Kakinuma, Hinako Shigihara for general technical support and Tetsuya Akiyama, Prof. Masashi Aoki for useful discussions (Tohoku University, Japan). Funding for this work was provided by Jiksak Bioengineering, Inc.

## AUTHOR CONTRIBUTIONS

Conceptualization, N.Y.

methodology, K.K.L.Y., J.K., D.I., and N.Y.

validation, K.K.L.Y., J.K., R.H and D.I.

formal analysis, K.K.L.Y., J.K., and D.I.

investigation, K.K.L.Y., J.K., D.I., and N.Y.

visualization, K.K.L.Y., J.K., and D.I.

writing – original draft, K.K.L.Y., J.K., and N.Y.

writing – review & editing, K.K.L.Y., J.K., D.I., N.S., and N.Y. Supervision, N.Y.

## DECLARATION OF GENERATIVE AI AND AI-ASSISTED TECHNOLOGIES IN THE WRITING PROCESS

During the preparation of this work, the author(s) used ChatGPT and Claude 3.5 Sonnet in order to improve on readability. After using this tool or service, the author(s) reviewed and edited the content as needed and take(s) full responsibility for the content of the publication.”

